# Phylogenomics reveals reticulate evolution in the *Chenopodium album* complex

**DOI:** 10.64898/2026.04.29.721649

**Authors:** Kate Escobar, Josefin Stiller, Pablo D. Cárdenas

## Abstract

**Background and Aims:** Complex genomic histories driven by hybridization and polyploidy can shape key plant traits such as defense, stress tolerance, and toxicity, particularly in Amaranthaceae, which includes crops such as quinoa and spinach. Within this family, white goosefoot (*Chenopodium album*) is both a widespread agricultural weed and a traditional food resource. However, its evolutionary history is complicated by discordant signals among genomic markers within the *C. album* complex, comprising diploid, tetraploid, and hexaploid taxa. Here, we tested whether reticulate evolution underlies this genome-wide discordance.

**Methods:** Using genome-scale phylogenomic data, we analysed 2,298 conserved nuclear loci (BUSCO genes) across 27 Amaranthaceae genomes. Both single- and multicopy gene families were included to capture signals of gene duplication, incomplete lineage sorting, and hybridization. Complementary phylogenomic approaches were used to evaluate whether the evolutionary history is best supported by strictly bifurcating relationships or by reticulate evolution.

**Key Results:** A consistent *C. album* lineage was recovered, comprising tetraploid and hexaploid *C. album* cytotypes together with *C. suecicum*, *C. strictum*, *C. formosanum*, *C. acuminatum*, and *C. opulifolium*. Phylogenetic discordance was concentrated within *Chenopodium*, particularly around the *C. album* and *C. quinoa* lineages. Models incorporating hybridization fit better than strictly bifurcating relationships, supporting at least two reticulation events. Hybridization signals were detected in 271 loci in tetraploid and 270 in hexaploid *C. album*, of which 232 were shared, indicating a shared hybrid origin rather than independent lineages.

**Conclusions:** The evolutionary history of the *C. album* lineage is best explained by reticulate processes involving hybridization and polyploidy. Conserved nuclear loci retain persistent signatures of these events, helping to resolve complex evolutionary histories in polyploid plant systems.

## Introduction

### Genome complexity and reticulate evolution in plants

Plant genomes exhibit structural and evolutionary complexity, shaped by recurrent processes such as whole-genome duplication (polyploidy), hybridization, and gene family expansion (Simpson 2019; Zuntini et al. 2024; McKain et al. 2018). Polyploidy has occurred repeatedly throughout angiosperm evolution and is widely recognized as a major driver of diversification and speciation (Baack and Rieseberg 2007; Madlung 2013). In particular, allopolyploidy combines distinct parental genomes within a single lineage, generating novel genetic combinations and phenotypic variation (Comai 2005; Mallet 2007; Rothfels 2021).

These processes reshape genome architecture by introducing multiple ancestral histories into a single genome, while gene duplication promotes functional divergence through subfunctionalization and neofunctionalization (Soltis et al. 2015; Guo et al. 2023; Zuntini et al. 2024). As a result, plant genomes often exhibit heterogeneous gene histories and reticulate patterns that challenge strictly bifurcating models of descent (Degnan and Rosenberg 2009; Stull et al. 2023). Signals of incomplete lineage sorting (ILS), hybridization, and gene duplication and loss (GDL) frequently act simultaneously, complicating phylogenetic inference and the interpretation of genome evolution (Maddison 1997; Yan et al. 2021; Guo et al. 2023; Steenwyk et al. 2023; Joyce et al. 2025).

### Ecological and agrifood relevance of Amaranthaceae

The family Amaranthaceae provides a compelling system in which to examine how these genomic processes shape ecological and agronomic traits (Fuentes-Bazan et al. 2012; Morales-Briones et al. 2021; Xu et al. 2024). This globally distributed lineage comprises approximately 2,000–2,500 species, including major crops such as quinoa (*Chenopodium quinoa*), spinach (*Spinacia oleracea*), sugar beet (*Beta vulgaris*), and grain amaranths (*Amaranthus* spp.) (Hernández-Ledesma et al. 2015; Jarvis et al. 2017). Many members exhibit strong tolerance to abiotic stress, including salinity, drought, and extreme temperatures, and are increasingly recognized as candidates for climate-resilient agriculture (Kadereit et al. 2012; Adolf et al. 2013; Marone et al. 2022; Sanfeliu Meliá and Cárdenas 2026). However, reconstructing evolutionary relationships within Amaranthaceae remains challenging due to extensive gene tree discordance arising from ILS, GDL, and hybridization (Sankoff 2018; Morales-Briones et al. 2021; Xu et al. 2024). These same processes are also expected to shape traits underlying ecological adaptation and agronomic value (Comai 2005; Soltis et al. 2015).

Within this context, *Chenopodium album* L. (white goosefoot) represents a particularly informative species. It is one of the most widely distributed angiosperms, occurring across temperate and subtropical regions on every inhabited continent (Williams 1963; Poonia and Upadhayay 2015; Tang et al. 2022; Shams et al. 2023) (Figure 1). Its success is closely associated with anthropogenic disturbance, and it functions as a highly competitive ruderal species characterized by rapid colonization, high fecundity, and efficient resource use (Bajwa et al. 2019; Krak et al. 2019). Consequently, *C. album* is widely regarded as a major agricultural weed, yet it has a long history of use as a leafy vegetable and pseudocereal in parts of Asia and Europe (Andreasen and Streibig 2011; Mueller-Bieniek et al. 2019; Kunwar et al. 2021).

**Figure 1.**
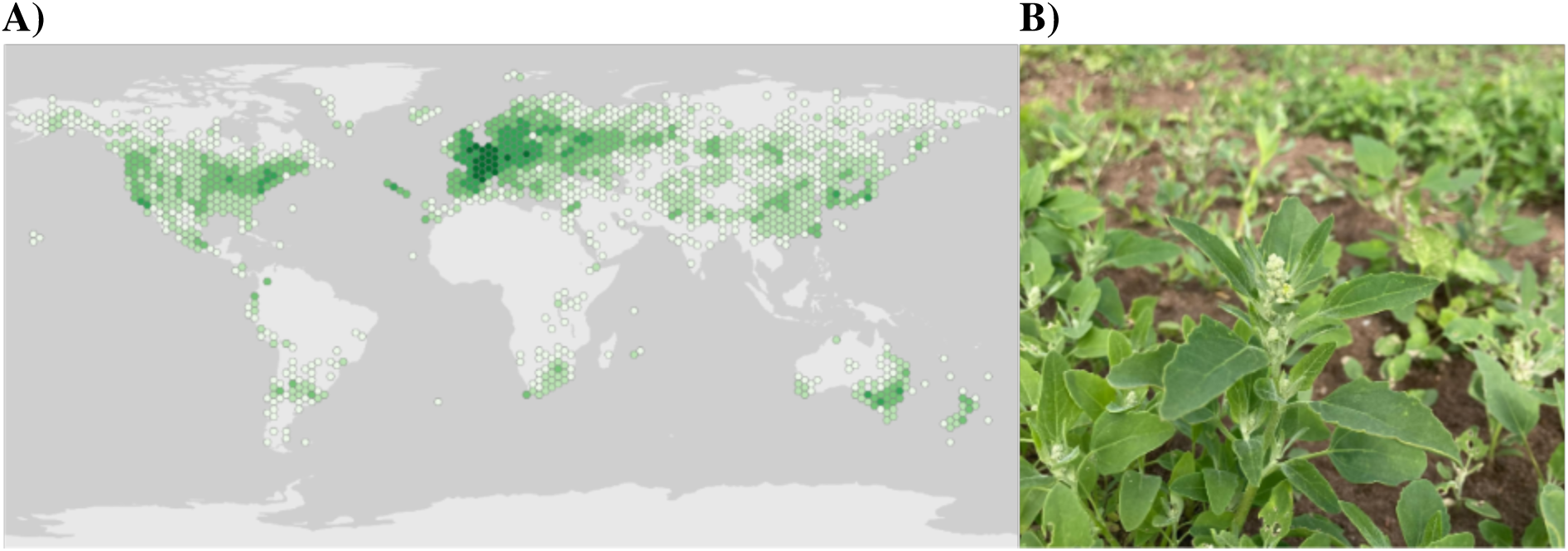
Global distribution and morphology of *Chenopodium album*. (A) Global occurrence records of *Chenopodium album* based on georeferenced observations from GBIF (GBIF 2023). Green hexagons indicate documented occurrences across the species’ distribution range, with darker shades representing higher occurrence density. (B) Vegetative morphology of *C. album*. Photo credit: Alexandra Sanfeliu Meliá.

### Specialized metabolites in plant defense and food use

Like many members of the Amaranthaceae, *C. album* produces specialized metabolites that mediate interactions with its environment, including defense against herbivores and pathogens and tolerance to abiotic stress (Pichersky and Gang 2000; Mueller-Bieniek et al. 2019; Weng et al. 2021; Song et al. 2022). Major classes, including phenolics and terpenoids such as saponins, are often lineage-specific and have diversified through gene duplication, subfunctionalization, and metabolic pathway evolution (Pichersky and Lewinsohn 2011; Huang and Dudareva 2023). The genes underlying these pathways exhibit diverse evolutionary trajectories, with some remaining conserved while others evolve rapidly through mutation, rearrangement, and neofunctionalization, often forming biosynthetic gene clusters characterized by high turnover (Chae et al. 2014; Soltis et al. 2015).

The dual role of specialized metabolites in enhancing plant fitness while limiting edibility creates a fundamental trade-off in crop development (D’Amelia et al. 2021; Huang and Dudareva 2023). This is well illustrated in quinoa, where domestication has targeted reduced bitterness while maintaining stress tolerance and agronomic performance (Patiranage et al. 2022; Wu et al. 2017; Guo et al. 2024; Cui et al. 2024; Guo et al. 2025). Triterpenoid saponins are major determinants of seed bitterness, with additional contributions from flavonoid glycosides in quinoa (Fiallos-Jurado et al. 2016; Suárez-Estrella et al. 2018; Pereira et al. 2020; Guo et al. 2024; Kollmar et al. 2025). Similar constraints are likely to apply to *C. album*, where evidence supports active saponin biosynthesis and its role in shaping chemical composition and associated traits (Lavaud et al. 2000; Fiallos-Jurado et al. 2016; Nedialkov and Kokanova-Nedialkova 2021; Vats et al. 2026). Interpreting variation in metabolic and agronomic traits therefore requires a genome-wide evolutionary framework that integrates signals across loci with different evolutionary histories and accounts for the reticulate evolution of the species.

### The *C. album* aggregate as a model of reticulate evolution

The *C. album* aggregate, as defined by Mandák et al. (2025), constitutes a polyploid species complex comprising closely related diploid, tetraploid, and hexaploid taxa connected through recurrent hybridization and polyploidization (Table 1). Within this framework, the aggregate is understood as an evolutionarily interconnected system of genome lineages (Mandák et al. 2025), with *C. album* sensu lato representing a focal lineage characterized by substantial cytotypic and genomic diversity (Mandák et al. 2016). Its evolutionary history has been investigated through approaches ranging from morphology and cytogenetics to molecular and genomic analyses, each contributing insight while also reflecting the limitations of available data and methods. Early taxonomic treatments relied primarily on morphological characters such as leaf shape, indumentum, and seed morphology, but their diagnostic value proved limited because of extensive phenotypic plasticity within the group (Linnaeus 1753; Moquin-Tandon 1840). This persistent ambiguity led the complex to be described as a “taxonomic receptacle” for morphologically variable forms (Wahl 1952).

**Table 1.** Conceptual definitions of terms used for the *C. album* complex.

| Term | Definition | Representative taxa | Genomic level |
| --- | --- | --- | --- |
| *C. album* aggregate (agg.) | Broad polyploid species complex comprising closely related diploid, tetraploid, and hexaploid taxa linked through recurrent hybridization and polyploidization. | "BB" lineage: *C. suecicum*, *C. ficifolium*, *C. ucrainicum*; "CCDD" lineage: *C. betaceum* (part of *C. strictum* complex), *C. glaucophyllum*, *C. novopokrovskyanum*, *C. striatiforme*; "BBCCDD" lineage: *C. album* s. str., *C. formosanum*, *C. giganteum*, *C. missouriense*, *C. perttii*, *C. probstii* | Multiple genome lineages and ploidy levels (Mandák et al. 2025). |
| *C. album* sensu lato (s. l.) | Broad taxonomic concept of *C. album*, referring to the morphologically variable focal lineage within the aggregate and encompassing multiple cytotypes. | Diploid, tetraploid and hexaploid *C. album* cytotypes | Cytotypically variable (Mehra and Malik 1963; Devi and Chrungoo 2015) |
| *C. album* sensu stricto (s. str.) | Narrower species concept corresponding to the hexaploid *C. album* lineage as treated in previous and recent genomic studies. | Hexaploid *C. album* | Defined allohexaploid genomic composition (BBC-CDD) (Mandák et al. 2025) |
| *C. album* lineage (this study) | Phylogenomic clade consistently recovered in this study, representing a subset of taxa within the broader *C. album* aggregate. | "*C. album* lineage": *C. opulifolium*, *C. acuminatum*, *C. strictum*, *C. formosanum*, *C. suecicum*, *C. album* (ploidy levels: 4x and 6x) | Mixed ploidy; includes diploid, tetraploid, and hexaploid representatives from the *C. album* aggregate |

The transition to cytogenetic approaches linked taxonomic uncertainty to underlying genome structure. Experimental crossing and chromosome counts enabled direct assessment of reproductive compatibility and ploidy levels to delimit lineages within the aggregate and showing that some putative taxa likely represented intraspecific variation rather than distinct species (Cole 1961). Later studies established *C. album* sensu stricto as predominantly hexaploid (n = 27; 2n = 54) (Uotila 1972), although additional cytotypes, including tetraploid and diploid forms, have also been reported within *C. album* sensu lato (Mehra and Malik 1963; Mandák et al. 2012; Devi and Chrungoo 2015; Krak et al. 2016; Habibi et al. 2018; Vats et al. 2026). Biochemical approaches further refined hypotheses of hybrid origin. Flavonoid profiling showed that diploid and tetraploid taxa possess distinct species-specific profiles, whereas hexaploid taxa exhibit additive or composite compositions derived from diploid species, particularly *C. suecicum* and *C. ficifolium*, supporting a hybrid origin and indicating retention of ancestral metabolic signatures following allopolyploidization (Rahiminejad and Gornall 2004).

The introduction of DNA sequence–based markers enabled more explicit reconstruction of evolutionary relationships. Plastid and nuclear ribosomal markers clarified higher-level relationships but often captured only a subset of the evolutionary history, typically reflecting a single parental lineage or a homogenized signal (Fuentes-Bazan et al. 2012; Viljoen et al. 2018; Park et al. 2021; Yao et al. 2019; Wei et al. 2023; Kiedaisch et al. 2025). Analyses of low-copy nuclear loci improved resolution. For example, the single-copy nuclear gene *SOS1* revealed distinct homeologous sequence clades corresponding to different ancestral genomes, supporting independent polyploid lineages across geographic regions (Walsh et al. 2015). Notably, reticulation is not solely a historical process, as ongoing homoploid hybridization between diploid taxa (*C. suecicum* × *C. ficifolium*) has been documented in Central Europe (Hodková and Mandák 2018).

Recent genome-wide and population-level analyses have provided a comprehensive framework for understanding these patterns. It et al. (2016) and Mandák et al. (2018) introduced the “genome-lineage” concept, in which multiple diploid lineages (A–H) act as recurring genomic building blocks across Eurasian polyploid taxa, allowing polyploid genomes to be partitioned into their constituent subgenomes. Building on this framework, Mandák et al. (2025) demonstrated that *C. album* sensu stricto is an allohexaploid with a BBCCDD genomic composition, comprising a diploid-derived BB subgenome and a CCDD component derived from an ancestral tetraploid lineage (Habibi et al. 2023). Importantly, these subgenomes do not correspond directly to extant species which could suggest that the parental lineages may be extinct or unsampled (Mandák et al. 2025). Multiple subgenome combination within *C. album* aggregate further support a polytopic origin involving repeated hybridization events (Mandák et al. 2025).

Biogeographic studies further contextualize this evolutionary complexity. Krak et al. (2019) identified Central Asia as a major centre of diversity for the *C. album* aggregate and linked its expansion into Europe to the spread of Neolithic agriculture, while Chrungoo et al. (2019) showed that Himalayan populations are closely related to *C. quinoa*-related lineages, highlighting regional differentiation and the influence of domestication-related processes on evolutionary trajectories. Mechanistic insights into genome evolution have also emerged from recent genomic studies. Belyayev et al. (2020*a*) demonstrated that satellite DNA evolution, often associated with transposable elements, plays a key role in genomic restructuring and stabilization following hybridization events (Belyayev et al. 2020*b*). More recently, high-resolution genomic analyses have implicated long terminal repeat (LTR) retrotransposon dynamics as an important driver of genome evolution within the complex (Jaggi et al. 2025), further emphasizing the role of repetitive elements in shaping polyploid genomes. Together, these findings show that the *C. album* aggregate represents a stabilized yet evolutionarily dynamic polyploid system.

However, these advances also highlight a fundamental limitation in current approaches to reconstructing the complex evolutionary history of the *C. album* aggregate (Smith et al. 2015; Smith and Hahn 2021). Analyses based on a limited number of loci, or restricted to single-copy genes to avoid issues of paralogy, exclude large portions of the genome that may contain important evolutionary signal (Smith and Hahn 2021; Zhang et al. 2021). Conversely, interpreting widespread gene tree discordance primarily as evidence of hybridization risks overestimating the extent of reticulation, as similar patterns can arise from ILS and GDL (Zhang et al. 2021; Stull et al. 2023). Distinguishing among these processes therefore remains a central challenge in plant phylogenomics.

### Disentangling genome-wide discordance using phylogenomics

Building on previous evidence for hybridization within the *C. album* aggregate, we implemented a genome-wide phylogenomic framework (Figure 2) to address three primary objectives. First, we quantified the extent of gene tree discordance to test whether patterns across the genome can be explained by ILS and GDL alone, or whether hybridization is required. Second, we evaluated whether two *C. album* cytotypes (4x and 6x) exhibit distinct patterns of genome-wide ancestry or share a common reticulate background. Third, we assessed whether extant species can be identified as potential progenitors of the polyploid genome.

**Figure 2.**
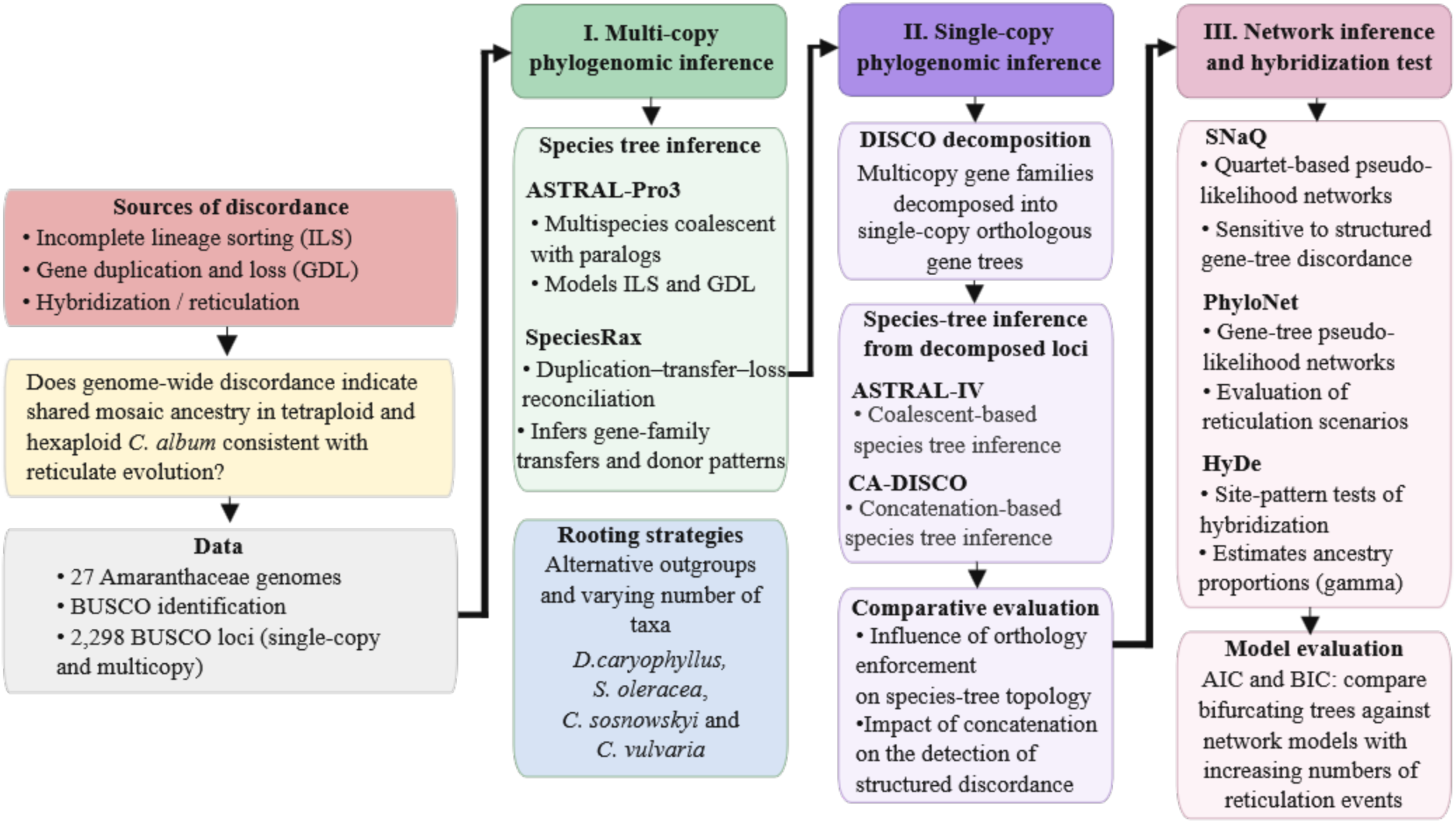
Phylogenomic framework for analysing genome-wide discordance in the *Chenopodium album* complex. Conserved BUSCO loci from sampled genomes were analysed using species tree inference, orthology-constrained approaches, and network and hybridization methods to evaluate bifurcating versus reticulate evolutionary models.

To achieve this, we analysed conserved BUSCO loci across representative Amaranthaceae genomes (Table 2). Rather than restricting analyses to single-copy genes, we explicitly incorporated both single- and multicopy loci to capture a broader proportion of the polyploid genome. This approach allowed us to model the combined effects of ILS, GDL and hybridization, thereby providing a comprehensive reconstruction of evolutionary relationships.

**Table 2.**
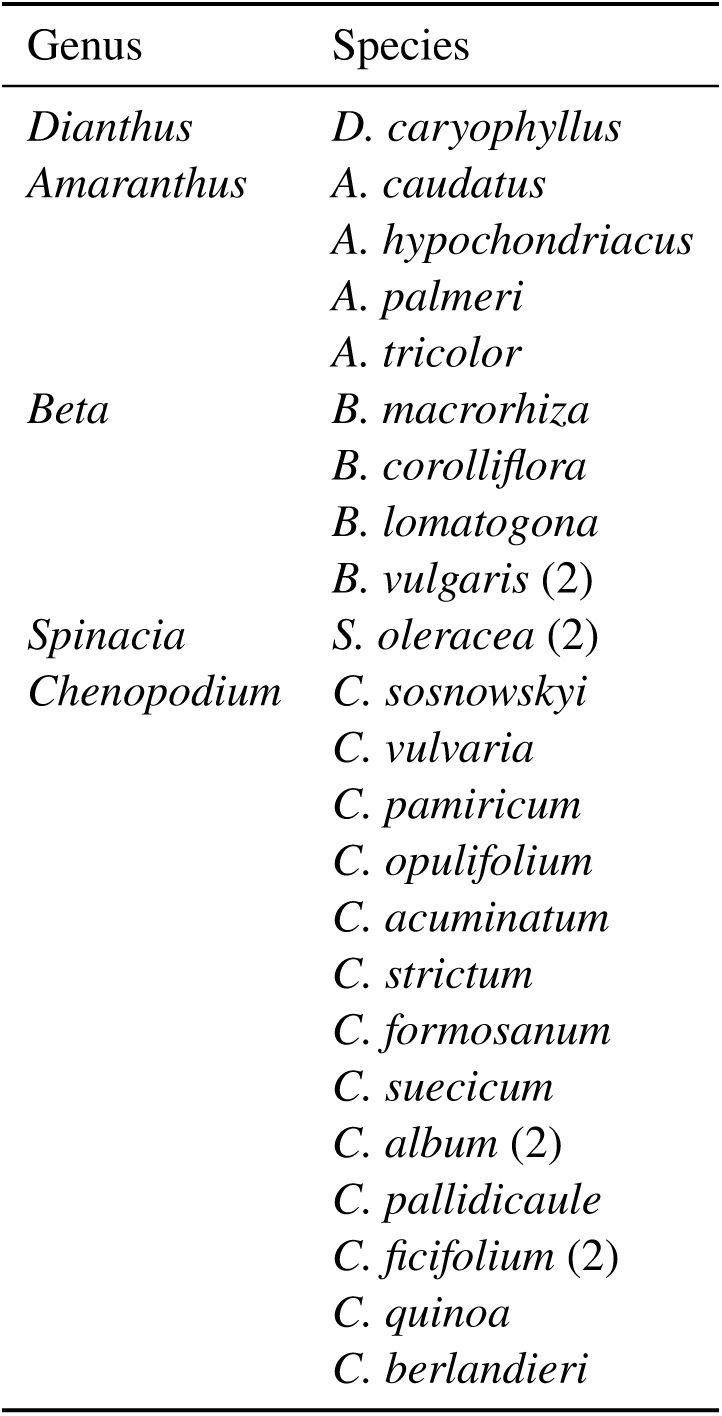
Taxa included in the phylogenomic analyses. Numbers in parentheses indicate the number of genome assemblies included per species.

## Materials and methods

### Phylogenomic data processing and gene tree inference

Genome assemblies representing 27 Amaranthaceae taxa (Table 2) were retrieved from the NCBI Genome database (Sayers et al. 2024) (Supplementary Table S1, Table S2, Figure S1). This sampling reflects currently available genomic resources and captures both diversity within the *Chenopodium* genus and broader phylogenetic representation across Amaranthaceae.

Orthologous loci were identified using BUSCO (v5.8.2), which stands for Benchmarking Universal Single-Copy Orthologs, with the *eudicots_odb10* dataset and MetaEuk gene prediction (Levy Karin et al. 2020; Manni et al. 2021*a*; Manni et al. 2021*b*). BUSCO loci are conserved protein-coding orthologs widely used as standardized markers of genome completeness and comparative genomics (Seppey et al. 2019). Both single-copy and multicopy BUSCO loci were retained to capture signals of ILS, GDL, and hybridization across the genome. To ensure data quality, only assemblies with at least 70% complete BUSCOs (≥1628 of 2326) were included in downstream analyses (Supplementary Table S3, Table S4).

For each BUSCO locus, amino acid sequences were aligned using MAFFT with the L-INS-i algorithm (Katoh et al. 2017). Codon-aware nucleotide alignments were generated by projecting aligned amino acids onto their corresponding coding sequences. Gene trees were inferred independently for each locus using maximum likelihood in IQ-TREE2 with automated model selection and 1000 ultrafast bootstrap replicates (Minh et al. 2020).

### Species tree inference using multicopy loci

Species trees were inferred using complementary frameworks designed to account for different sources of gene tree discordance. First, ASTRAL-Pro3 was applied to multicopy gene trees to infer a coalescent-based species tree while explicitly modelling incomplete lineage sorting (ILS) and gene duplication and loss (GDL) (Zhang et al. 2025). This approach identifies speciation-driven quartets within gene family trees, allowing phylogenetic signal from multicopy loci while reducing the impact of paralogy-driven conflict (Zhang et al. 2021; Zhang and Mirarab 2022). Branch lengths were estimated in coalescent units, such that internal branch lengths reflect the amount of genealogical separation in the multispecies coalescent framework (Degnan and Rosenberg 2009; Jiang et al. 2020). Quartet support values (q_1_, q_2_, and q_3_) were used to summarize localized gene tree discordance across the inferred species tree (Zhang et al. 2025).

Second, SpeciesRax was used to infer a species tree under a maximum-likelihood framework that models gene family evolution through duplication, transfer, and loss (DTL) (Morel et al. 2022). In addition to species tree estimation, SpeciesRax infers gene transfer events by reconciling gene trees with the species tree under the UndatedDTL model, identifying the most likely donor and recipient lineages (Morel et al. 2020; Morel et al. 2022). The inferred transferred events are most parsimoniously interpreted as gene tree–species tree incongruence rather than literal horizontal gene transfer (Morel et al. 2022). Furthermore, this framework does not explicitly model ILS, so inferred transfer events were interpreted conservatively as signatures of gene tree–species tree incongruence and evaluated alongside coalescent-based results.

To assess the robustness of species tree inference to outgroup selection, analyses were repeated using alternative outgroups, including *Dianthus caryophyllus*, *Spinacia oleracea*, and a combined outgroup of *C. sosnowskyi* and *C. vulvaria*. These comparisons were used to evaluate the impact of rooting on topology and branch support.

### Species tree inference using single-copy loci

Several downstream analyses required orthologous (single-copy) gene trees. Therefore, multicopy BUSCO trees were decomposed using DISCO (Decomposition Into Single-Copy Orthologs) to extract putative orthologous subtrees (Willson et al. 2021). Resulting trees were filtered to retain loci with one copy per species and at least four taxa.

These orthology-constrained datasets were used for species tree estimation with ASTRAL-IV (v1.24.4.8) (Mirarab et al. 2014; Zhang et al. 2025) and for concatenation-based analyses, allowing comparison between multicopy-aware and orthology-restricted inference frameworks (Yu et al. 2014; Minh et al. 2020; Jiang et al. 2020; Zhang et al. 2025). Species trees inferred with ASTRAL-IV were annotated with quartet support values and branch lengths in coalescent units.

### Phylogenetic network inference

Phylogenetic networks were inferred using two pseudolikelihood-based approaches. First, PhyloNet (v3.8.4) was used to infer gene-tree-based networks under a maximum pseudolikelihood framework (Wen et al. 2018). Analyses were conducted on rooted gene trees derived from DISCO-decomposed BUSCO loci for 15 *Chenopodium* taxa, with *S. oleracea* specified as the outgroup. Because PhyloNet does not explicitly model gene duplication and loss, analyses were restricted to single-copy gene trees to reduce potential biases associated with paralogy (Altenhoff et al. 2019; Yan et al. 2021). Models allowing 0–3 reticulations were evaluated using multiple replicate searches, with branch lengths and inheritance probabilities optimized during and after inference. Second, network inference was performed using SNaQ within PhyloNetworks (v1.1.0), which estimates networks by maximizing a quartet-based pseudolikelihood derived from gene tree concordance factors (Solís-Lemus and Ané 2016; Solís-Lemus et al. 2017). Analyses were based on the same DISCO-decomposed gene trees. Concordance factors were estimated from gene trees, and network searches were conducted for ℎ = 0–3, with replicate runs initialized from the ASTRAL-IV species tree.

For both approaches, model fit was evaluated using pseudolikelihood scores and compared across reticulation levels using Akaike (AIC) and Bayesian (BIC) information criteria, calculated as AIC = 2k − 2 ln L and BIC = k ln n − 2 ln L, where k is the number of free parameters, n is the number of gene trees, and ln L is the maximum pseudolikelihood score. Because pseudo-likelihoods are approximations of full likelihoods, AIC and BIC were interpreted comparatively to assess relative support among models with increasing reticulation complexity (Akaike 1974; Schwarz 1978; Susko and Roger 2019).

### Hybridization testing

Hybridization was further evaluated using HyDe (Hybridization Detection using Phylogenetic Invariants), which tests for admixture among taxon triplets based on site pattern frequencies (Blischak et al. 2018). Analyses were conducted on BUSCO alignments by testing all combinations of ingroup taxa, with *S. oleracea* specified as a fixed outgroup. Each test evaluated a triplet in the form (P1, Hybrid, P2) relative to the outgroup.

Bootstrap resampling (1000 replicates) was performed across loci to estimate ancestry proportions (γ) and assess statistical support. Locus-level summaries were obtained by collapsing bootstrap replicates, and loci with ≥ 95% significant replicates were considered strongly supported. Folded γ̂ values were used to quantify hybridization, where γ̂ represents the proportion of ancestry inherited from parental lineage P1, and the frequency of inferred parental combinations was summarized across loci to characterize genome-wide patterns of admixture (Martin et al. 2014; Blischak et al. 2018; Kubatko and Chifman 2019).

Unlike network-based approaches, HyDe does not reconstruct explicit phylogenetic networks but instead provides locus-level evidence for hybridization (Blischak et al. 2018; Kubatko and Chifman 2019). Candidate introgressed loci identified from HyDe analyses were functionally annotated using orthology-based assignments from BUSCO and OrthoDB (Manni et al. 2021*a*; Zdobnov et al. 2021). Introgressed loci were further classified as shared between cytotypes or unique to each cytotype based on their occurrence across the tetraploid and hexaploid *C. album* genomes, and their functional annotations were used to summarize the broad categories of genes represented.

## Results

### BUSCO duplication patterns reflect ploidy across Amaranthaceae genomes

Genome assemblies exhibited variation in contiguity metrics (i.e., contig and scaffold N50; Supplementary Figure S2, Figure S3), yet BUSCO analyses indicated consistently high completeness across most taxa (Figure 3). Fragmented BUSCOs ranged from 0.2% to 0.8%, and missing BUSCOs were generally low (1.4%–3.5%), with the exception of *B. macrorhiza* (13.8% missing; 14.6% fragmented) (Supplementary Table S3).

**Figure 3.**
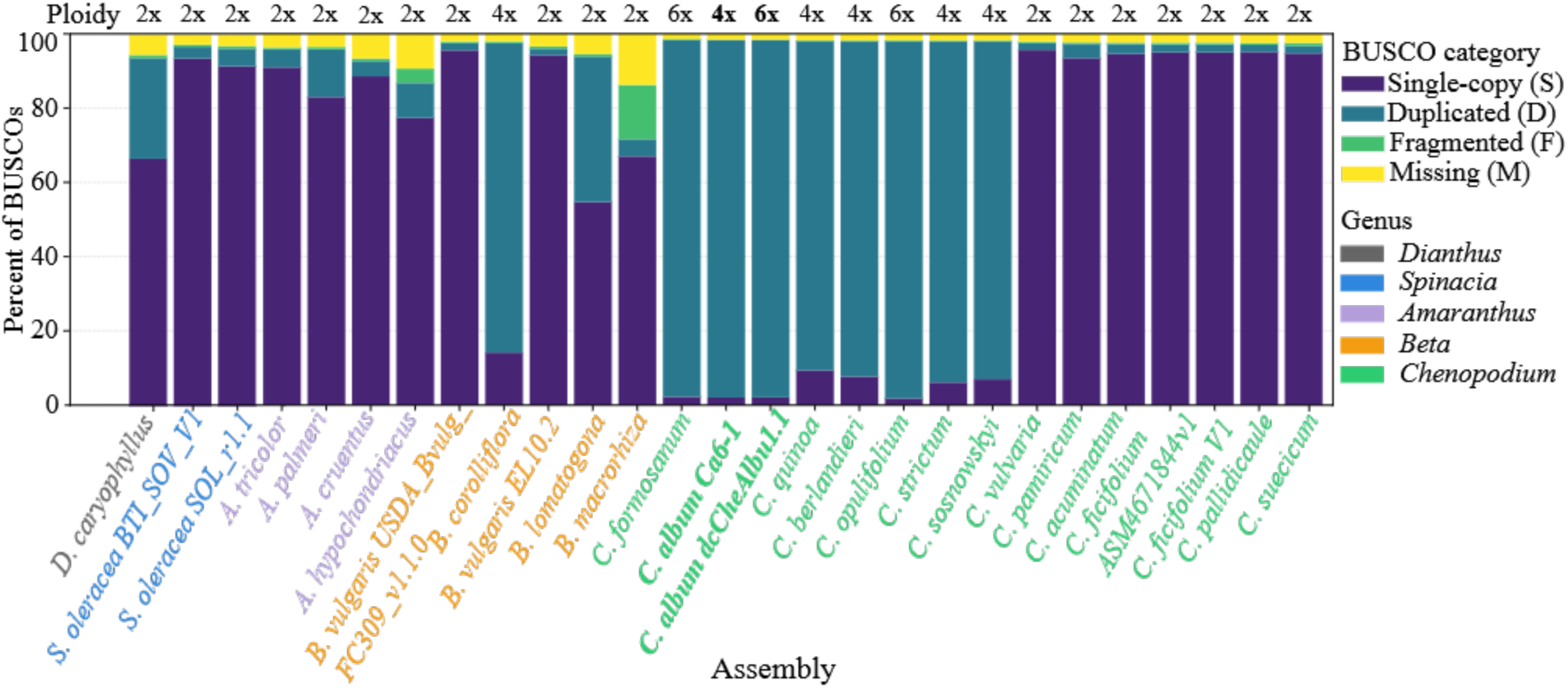
BUSCO composition across Amaranthaceae genome assemblies. Stacked bars show the proportion of BUSCO genes classified as complete single-copy (S), complete duplicated (D), fragmented (F), or missing (M) using the *eudicots_odb10* dataset. Overall BUSCO completeness corresponds to the combined proportion of single-copy and duplicated genes (C = S + D), which is consistently high across most assemblies. Polyploid *Chenopodium* genomes are enriched in duplicated BUSCOs, whereas diploid taxa are dominated by single-copy BUSCOs. The low proportion of fragmented and missing BUSCOs further supports the suitability of these assemblies for phylogenomic analyses.

In contrast to relatively uniform completeness, BUSCO duplication patterns varied strongly among taxa and closely tracked known ploidy levels. Polyploid *Chenopodium* genomes were dominated by duplicated BUSCOs, with duplication proportions ranging from 88.7% in *C. quinoa* to 96.2% in both tetraploid and hexaploid *C. album* assemblies, and 96.1% in *C. formosanum*. Similarly high duplication levels were observed in other polyploids, including *C. opulifolium* (96.2%), *C. sosnowskyi* (91.0%), *C. strictum* (91.9%), and *C. berlandieri* (90.3%).

Diploid *Chenopodium* species were characterized by predominantly single-copy BUSCOs (93.5%–95.7%) and low duplication levels (1.8%–3.8%), consistent with expectations for non-polyploid genomes. Similar patterns were observed in other diploid Amaranthaceae taxa, including *S. oleracea* (3.1%–4.7% duplicated) and *B. vulgaris* (1.7%–1.9% duplicated). Two diploid taxa deviated from this trend: *B. lomatogona* exhibited substantially higher duplication levels (39.2%), and *D. caryophyllus* also showed elevated duplication (27.1%). This may reflect lineage-specific genome duplication or assembly artifacts.

Overall, BUSCO duplication profiles closely matched known ploidy levels and provide independent genomic evidence for extensive gene duplication in polyploid *Chenopodium*. These patterns highlighted the prevalence of multicopy gene families in the dataset and motivated the use of duplication-aware phylogenomic approaches in downstream analyses.

### Phylogenetic discordance is concentrated in the *C. album* and *C. quinoa* lineages

Species trees inferred using the multicopy-aware framework ASTRAL-Pro3 recovered congruent higher-level relationships across Amaranthaceae (Figure 4). Outgroup choice did not affect relationships within *Chenopodium* (Figure 5), and subsequent analyses were therefore rooted using *D. caryophyllus*. Despite differing model assumptions, ASTRAL-Pro3 (accounting for ILS and GDL) and SpeciesRax (modelling duplication–transfer–loss) both consistently recovered monophyly of *Chenopodium* and similar relationships among major lineages (Supplementary Figure S4).

**Figure 4.**
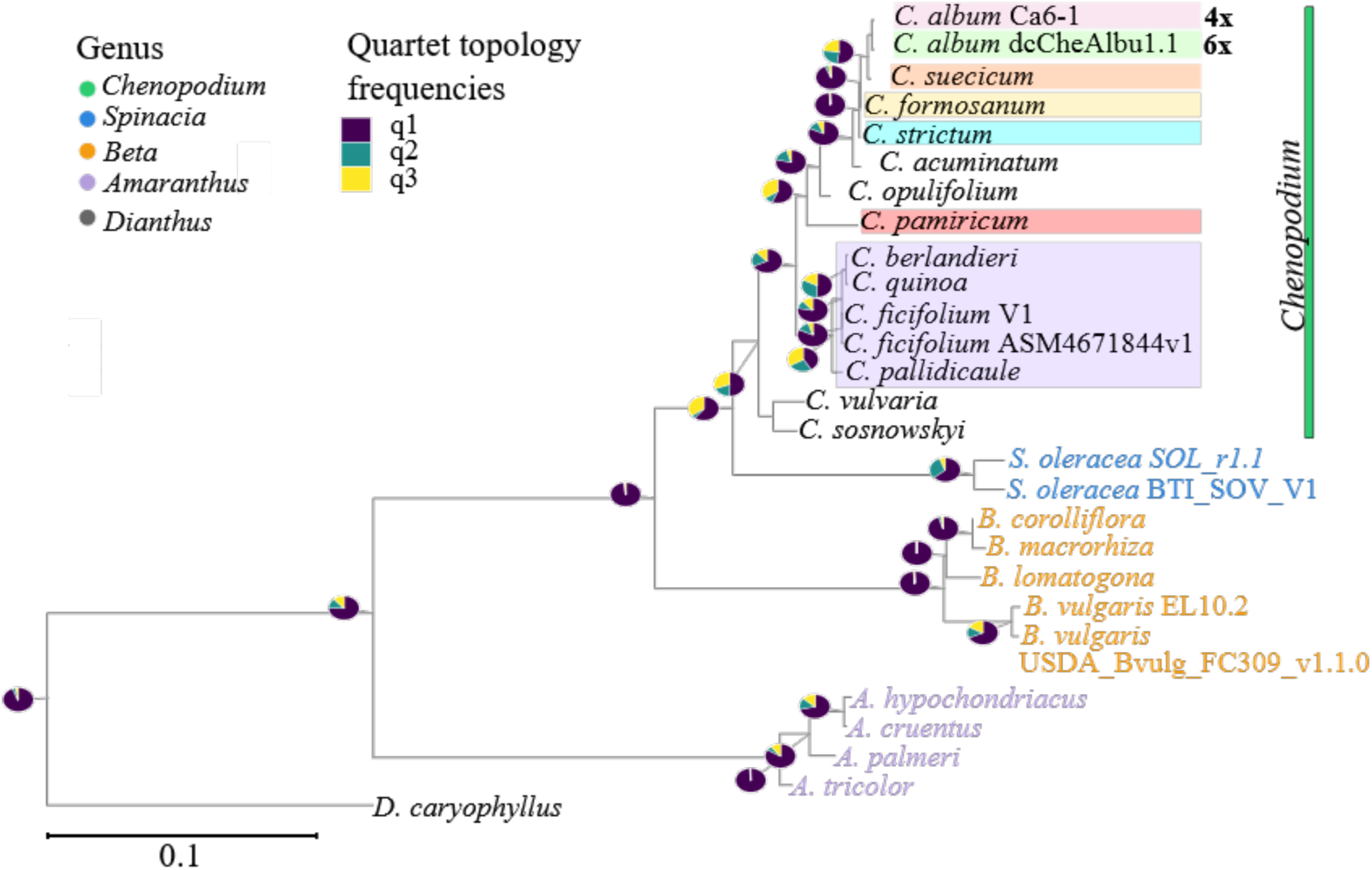
Species tree inferred with ASTRAL-Pro3 from 2,292 BUSCO gene family trees across 27 taxa. *D. caryophyllus* was used as outgroup. Branch lengths are shown in coalescent units. Pie charts at nodes indicate the relative frequencies of the three possible quartet topologies, where q1 matches the displayed species-tree relationship and q2–q3 represent the two alternative topologies.

**Figure 5.**
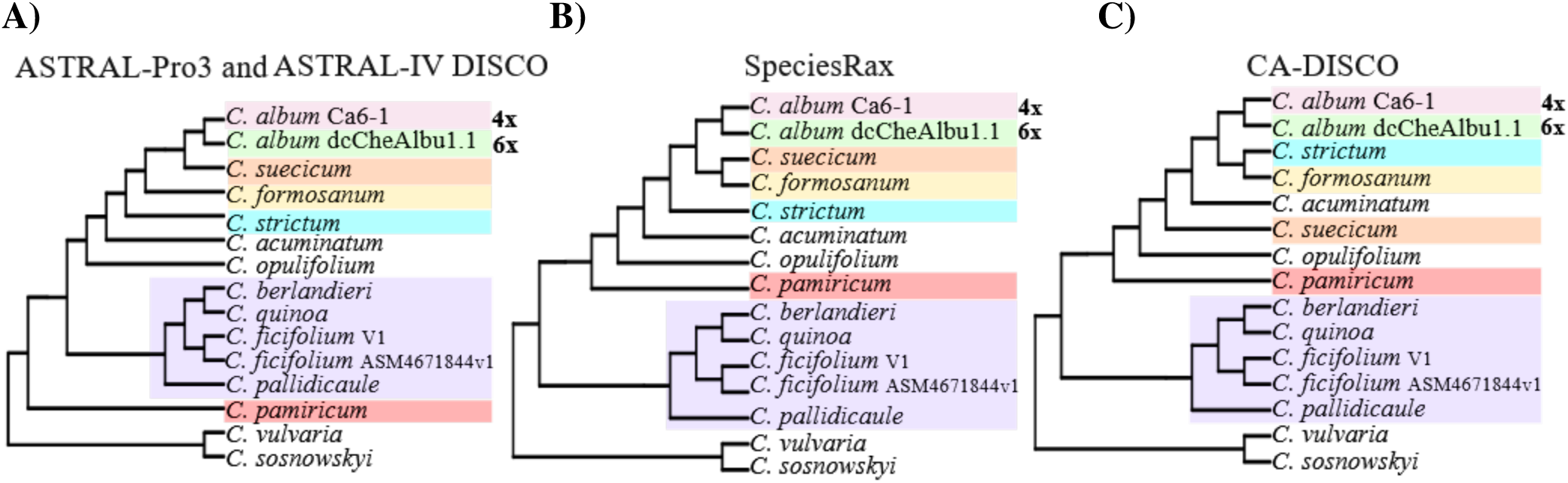
Alternative species-tree topologies inferred for *Chenopodium*. Species trees inferred using (A) ASTRAL-Pro3/ASTRAL-IV (DISCO), (B) SpeciesRax, and (C) concatenation (CA-DISCO). The core *C. album* lineage (highlighted) is consistently recovered across methods, whereas relationships within the quinoa-associated clade and among other *Chenopodium* taxa vary among analyses, indicating localized gene-tree discordance.

Across analyses, a *C. album* lineage comprising the two *C. album* cytotypes (Ca6-1 (4x) and dcCheAlbu1.1 (6x)), together with *C. suecicum*, *C. formosanum*, *C. strictum*, *C. acuminatum*, and *C. opulifolium*, was consistently recovered. Similarly, relationships within the *C. quinoa* lineage (*C. quinoa*, *C. berlandieri*, *C. ficifolium*, and *C. pallidicaule*) were stable across methods (Supplementary Figure S6 - Figure S11). These results indicate that both clades are robust to inference framework and outgroup selection.

Quartet support values from ASTRAL-Pro3 (Figure 4) indicate substantial gene tree discordance within *Chenopodium*. While many deeper nodes are dominated by a single topology (high q_1_), several internal branches show support for alternative quartet resolutions. Conflicting signal is evident not only within *Chenopodium*, but also in parts of *Amaranthus*, indicating discordance across multiple levels of the tree. Within *Chenopodium*, however, discordance is particularly pronounced in the region containing the *C. album* and *C. quinoa*-associated lineages. Specifically, the placement of *C. pamiricum* shows strong conflict and varies across inference methods (Figure 5). Similarly, high discordance is evident at the node uniting *C. suecicum* with the two *C. album* cytotypes. Nodes within the *C. quinoa*-associated clade, including relationships among *C. quinoa*, *C. berlandieri*, *C. ficifolium*, and *C. pallidicaule*, also show substantial alternative quartet support, consistent with heterogeneous signal across loci.

SpeciesRax reconciliation analyses further supported discordance in the same region with substantial numbers of gene transfer events involving *C. album* lineages (Supplementary Figure S5). In *C. album* dcCheAlbu1.1 (6x), the highest number of inferred transfers was associated with the internal node shared with *C. formosanum* (551), rather than with *C. formosanum* itself. Additional inferred transfer events were associated with the internal node shared with *C. strictum* (230), as well as with extant taxa, including *C. strictum* (158) and *C. suecicum* (137). A similar pattern was evident for *C. album* Ca6-1 (4x), where the majority of inferred transfers were associated with internal nodes within the same clade, particularly those related to *C. suecicum* (370) and *C. strictum* (194). In contrast, *C. quinoa* exhibited substantially fewer inferred transfer events overall, with the highest contributions arising from internal nodes within the *C. album* lineage, including the node shared with *C. suecicum* (15) and the node uniting the tetraploid and hexaploid *C. album* cytotypes (14). Additional inferred transfers were associated with extant taxa, including *C. formosanum* (12) and *C. opulifolium* (11).

To assess the effect of restricting analyses to putative orthologous loci, species trees were also inferred from DISCO-decomposed gene families. ASTRAL-IV analyses of these loci recovered a topology largely consistent with multicopy-aware methods, again supporting the *C. album* lineage (Figure 5). In contrast, concatenation-based inference (CA-DISCO) produced an alternative arrangement within the *C. album*-associated clade.

### Phylogenetic network analyses support at least two hybridization events in *Chenopodium*

Both PhyloNet and SNaQ recovered a consistent pattern in which model fit improved strongly with the addition of one and two reticulation events, followed by little or no further improvement (Tables 3, 4). This pattern indicates that most of the signal for reticulation is captured by approximately two hybridization events.

**Table 3.** Model comparison for PhyloNet network inference across increasing numbers of reticulations (ℎ). Values correspond to the best replicate for each ℎ, where ℎ = 3 (in bold) showed lowest AIC and BIC values and k is the number of free parameters.

| $h$ | $\ln L_{\text{pseudo}}$ | $k$ | AIC | BIC |
| --- | --- | --- | --- | --- |
| 0 | -242865.98 | 15 | 485762.0 | 485848.1 |
| 1 | -239188.57 | 17 | 478411.1 | 478508.7 |
| 2 | -235706.02 | 19 | 471450.0 | 471559.1 |
| 3 | **-235243.59** | 21 | **470529.2** | **470649.7** |

**Table 4.** Model comparison for SNaQ network inference across increasing numbers of reticulations (ℎ). Values correspond to the best replicate for each ℎ, where ℎ = 2 (in bold) showed lowest AIC and BIC values and k is the number of free parameters.

| $h$ | $\ln L_{\text{pseudo}}$ | $k$ | AIC | BIC |
| --- | --- | --- | --- | --- |
| 0 | -25744.5 | 15 | -51458.9 | -51372.8 |
| 1 | -24857.6 | 17 | -49681.1 | -49583.6 |
| 2 | **-17436.0** | 19 | **-34834.1** | **-34725.0** |
| 3 | -19767.8 | 21 | -39493.7 | -39373.2 |

In PhyloNet, pseudolikelihood increased substantially from ℎ = 0 to ℎ = 2, with only a minor additional gain at ℎ = 3 (Table 3) (Supplementary Table S5, Table S6 and Figure S12. While AIC continued to favor more complex models, BIC showed only marginal improvement beyond ℎ = 2, consistent with a plateau in model fit. SNaQ analyses showed a similar trajectory, with model fit improving up to ℎ = 2 but declining at ℎ = 3, and both AIC and BIC supported ℎ = 2 as the best model (Table 4) (Supplementary Table S7, Figure S12).

Despite this agreement in model complexity, the inferred network topologies differ between methods (Figure 6) (Supplementary Figure S12). Both PhyloNet and SNaQ recover the major *C. album* and *C. quinoa* lineages; however, internal relationships within the *C. album* complex are not consistent. In PhyloNet, the ℎ = 2 and ℎ = 3 networks place the tetraploid *C. album* with *C. suecicum*, while the hexaploid cytotype is recovered separately, resulting in a non-monophyletic arrangement of *C. album* s. l.. In contrast, SNaQ networks at comparable reticulation levels recover the tetraploid and hexaploid cytotypes grouped together.

**Figure 6.**
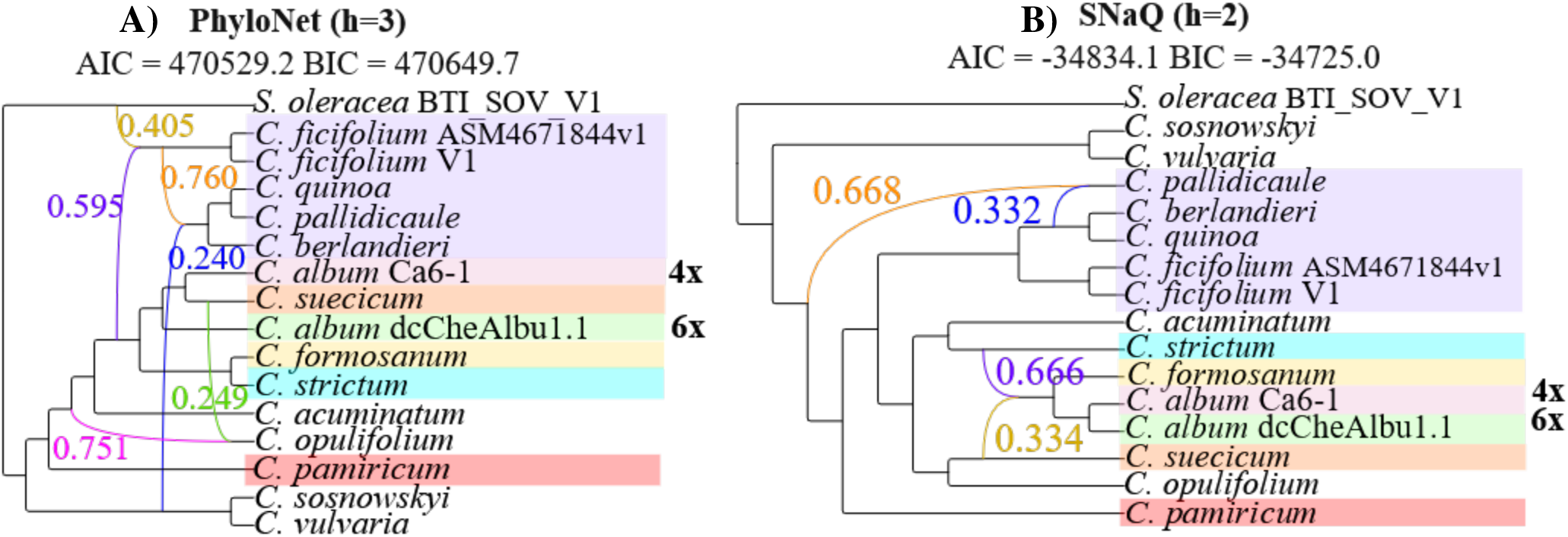
Comparison of phylogenetic network inference methods. Inferred networks through (A) PhyloNet and (B) SNaQ. Coloured edges indicate inferred hybridization events, with inheritance probabilities (γ), where each pair of edges, in complementary colours, sums up to 1. Both methods support models with approximately 2 reticulation events, but differ in the placement of relationships within the *C. album* complex.

Although both methods support a similar number of reticulation events, relationships within the *C. album* lineage remain method-dependent. Taken together, these results indicate that at least two hybridization events are required to explain the evolutionary history of the *C. album* lineage, while additional reticulations provide limited improvement in model fit and do not consistently resolve relationships among cytotypes.

### HyDe analyses reveal shared but heterogeneous signals of hybridization across BUSCO loci

HyDe analyses were conducted independently for the tetraploid (*C. album* Ca6-1) and hexaploid (*C. album* dcCheAlbu1.1) cytotypes across 2,298 BUSCO loci. A locus was considered to support hybridization when ≥ 95% of bootstrap replicates were significant (p < 0.05) . Under this criterion, 271 loci supported hybridization in Ca6-1 (4x) and 270 loci in dcCheAlbu1.1 (6x), corresponding to approximately 13.5% of all loci. Of these, 232 loci were shared between cytotypes, whereas 39 and 38 loci were unique to Ca6-1 (4x) and dcCheAlbu1.1 (6x), respectively. This substantial overlap indicates that most locus-level hybridization signals are shared between cytotypes.

The distribution of inheritance proportion estimates (γ̂) was similar between cytotypes (Figure 7). In both cases, γ̂ values showed a high peak around 0.5, with fewer loci exhibiting values close to 0 or 1. This pattern indicates that many loci provide limited resolution among alternative parental configurations, while a smaller subset supports more asymmetric ancestry. Notably, the total counts differ by approximately one order of magnitude between cytotypes, likely reflecting differences in the number of informative sites associated with differences in ploidy level.

**Figure 7.**
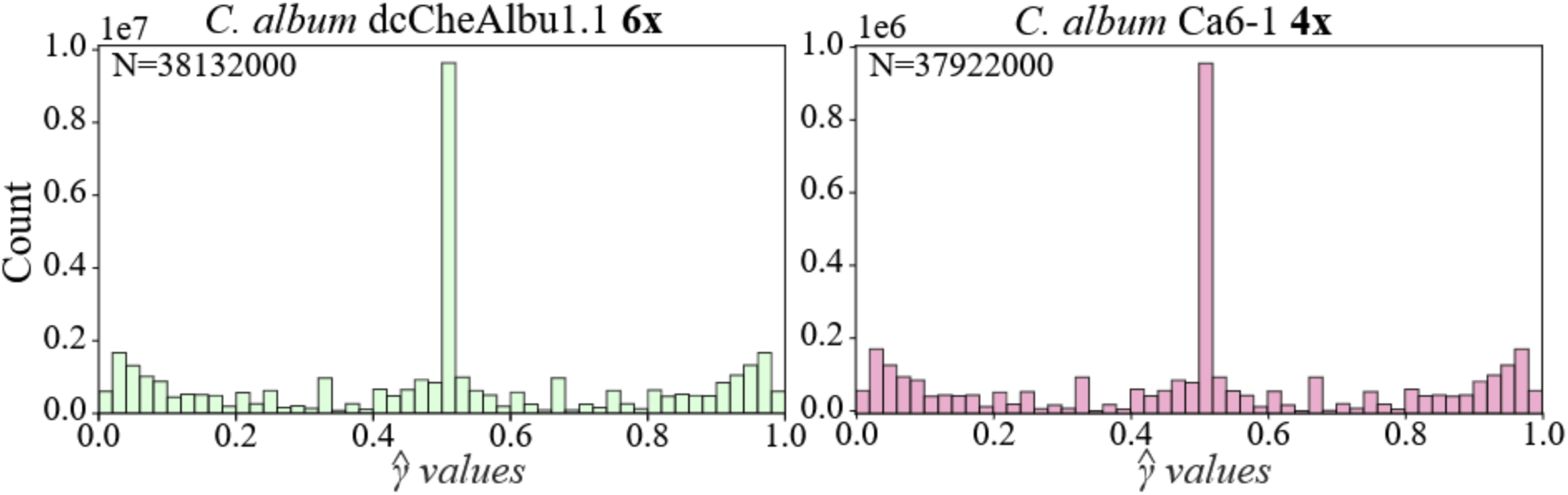
Distribution of inheritance estimates (γ̂) across the two *C. album* cytotypes. Histograms show the distribution of γ̂ values from HyDe analyses (1000 bootstrap replicates) for hexaploid and tetraploid *C. album*. Values near 0.5 indicate limited resolution among alternative parental configurations, whereas values closer to 0 or 1 indicate increasingly asymmetric support for specific parental relationships. N denotes the total number of bootstrap γ̂ estimates contributing to each histogram (across BUSCO loci, triplets, and replicates).

Across loci meeting the support threshold, *C. acuminatum*, *C. opulifolium*, and *C. pamiricum* were most frequently involved in supported triplets for both cytotypes (Figure 8). The ranking of contributing taxa was broadly consistent between dcCheAlbu1.1 (6x) and Ca6-1 (4x), indicating that similar lineages are repeatedly associated with hybridization signals. To facilitate interpretation, these ranked genomic contributors were also visualized geographically, with shading intensity reflecting their relative contribution to each focal *C. album* cytotype (Supplementary Figure S13, Figure S14). These results show that hybridization signals are distributed across a subset of loci and involve a similar set of taxa in both cytotypes. Together, this overlap suggests that tetraploid and hexaploid *C. album* cytotypes derive from a shared reticulate ancestry rather than independent hybridization events.

**Figure 8.**
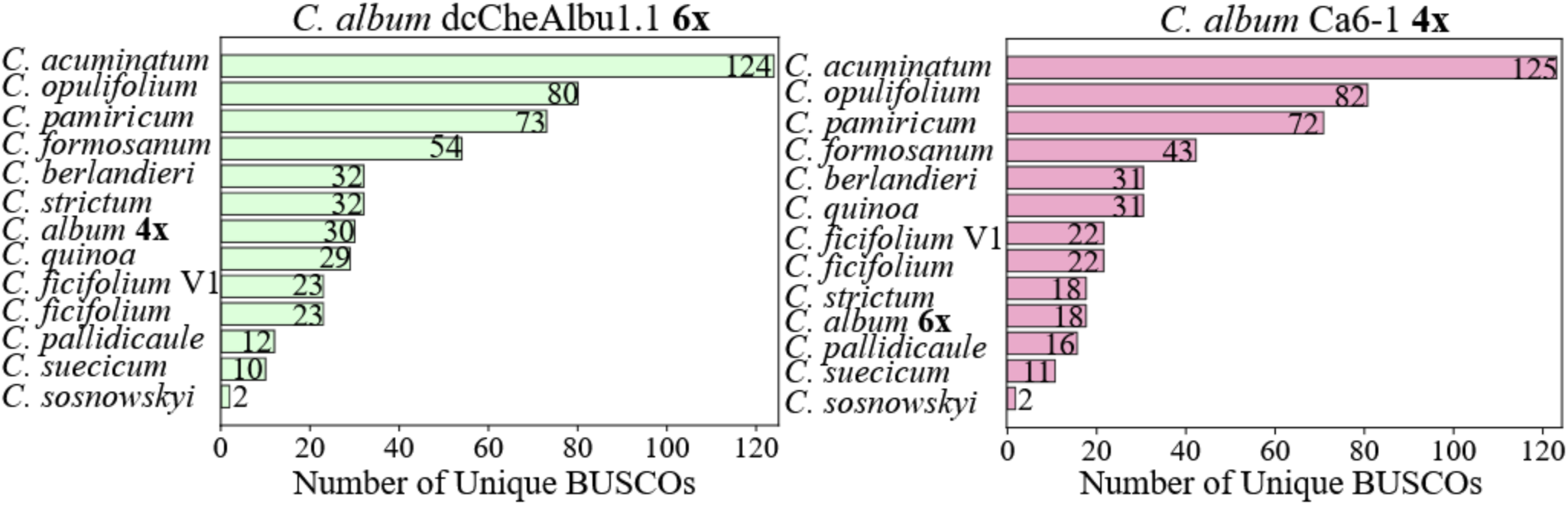
BUSCO loci supporting parental relationships in HyDe analyses. Bar plots show the number of loci in which each taxon was recovered as a parental contributor in significantly supported triplets for *C. album* dcCheAlbu1.1 (6x) and Ca6-1 (4x). Only loci meeting the support threshold (≥ 95% significant bootstrap replicates) are included.

### Introgressed loci across *C. album* cytotypes span broad functional categories

BUSCO loci identified as introgressed in both *C. album* cytotypes correspond to conserved orthologous groups spanning a wide range of fundamental cellular processes (Supplementary Table S8). Homology-based annotation indicates that these loci include genes associated with genome maintenance (e.g., DNA repair and replication), gene regulation (e.g., chromatin-associated and transcription-related proteins), RNA processing and translation, organelle-associated functions, intracellular trafficking, post-translational modification, and general metabolic processes. Cytotype-specific introgressed loci show similar broad functional profiles, without clear restriction to particular biological pathways (Supplementary Table S9, Table S10). Because BUSCO loci represent highly conserved orthologs, these annotations are expected to reflect general cellular functions. Accordingly, these results provide a qualitative overview of the functional diversity represented among introgressed loci, rather than evidence for functional enrichment or cytotype-specific adaptation.

## Discussion

### Reticulate evolution and taxonomic ambiguity in the *C. album* complex

Our analyses indicate that the evolutionary history of the *C. album* lineage is better explained by reticulate processes than by a strictly bifurcating model of divergence. Across multiple inference frameworks, phylogenetic conflict was concentrated within the *C. album* and *C. quinoa* lineages, whereas deeper relationships across Amaranthaceae were comparatively stable. Phylogenetic network analyses further supported models incorporating hybridization, with two reticulation events capturing most of the signal. At the locus level, HyDe analyses revealed extensive overlap in hybridization signals between tetraploid and hexaploid *C. album* cytotypes, with 232 shared loci compared to only 38–39 cytotype-specific loci. Together, these results indicate that both cytotypes derive from a shared reticulate genomic background shaped by historical hybridization and polyploidization, rather than representing independent recent hybrid origins. The broad functional distribution of introgressed BUSCO loci further suggests that these processes have affected the genome widely, consistent with genomic restructuring following hybridization and whole-genome duplication.

These findings provide a mechanistic explanation for the long-standing taxonomic ambiguity associated with the *C. album* complex. Historically, variation in morphology and ploidy led to contrasting classifications in which diploid, tetraploid, and hexaploid forms were treated either as distinct taxa or as intraspecific variants. However, the consistent recovery of a broader *C. album* lineage encompassing taxa such as *C. suecicum*, *C. strictum*, *C. formosanum*, *C. opulifolium*, and *C. acuminatum* suggests that these entities are not fully discrete evolutionary units, but components of an interconnected reticulate complex of closely related lineages. In this context, conflict among morphological, cytogenetic, and phylogenomic classifications is expected because each framework captures different aspects of a complex reticulate system. The *C. album* aggregate is therefore best understood as an evolutionarily interconnected polyploid complex, within which the *C. album* lineage identified here represents a phylogenomically coherent unit. This interpretation reconciles historical taxonomic treatments with genomic evidence and highlights the importance of explicitly accounting for reticulate evolution when delimiting species boundaries in polyploid plant systems.

Beyond its taxonomic implications, the *C. album* complex, alongside the *C. quinoa* lineage, provides a useful model system for studying reticulate evolution in plants. Its combination of multiple ploidy levels, recurrent hybridization, and partially stabilized genomic architectures provides an opportunity to investigate how hybridization and polyploidization shape genome evolution over time. At the same time, this complexity underscores a key methodological challenge: different markers and analytical approaches may recover distinct, and sometimes conflicting, evolutionary signals. As shown here, these patterns of discordance are not simply analytical artefacts, but reflect the underlying reticulate history of the group and are not fully captured by strictly bifurcating models. This highlights the value of using both single-copy and multicopy genes, complementing tree-based approaches with phylogenetic network inference and locus-level tests of hybridization.

Finally, the reticulate and polytopic origins of *C. album* may also contribute to its ecological success. Genomic evidence indicates that *C. album* s. str. originated repeatedly from distinct ancestral lineages, generating mosaic subgenomic combinations associated with substantial morphological and ecological plasticity (Mandák et al. 2025). The preservation of ancestral variation within its subgenomes, together with repeated hybridization events, may enhance adaptive potential by combining diverse genetic backgrounds within a single evolutionary lineage. In this context, the global success of *C. album* as a widespread and highly adaptable weed may reflect not only its polyploid nature, but also genomic plasticity within a reticulate framework.

### Linking phylogenomic frameworks to specialized metabolism

The results presented highlight the potential of using conserved, genome-wide nuclear loci for reconstructing evolutionary history in complex polyploid systems. Because BUSCO genes represent broadly conserved orthologs, they provide a relatively stable phylogenetic framework that is less influenced by rapid sequence divergence, and are therefore suited to resolving deep shared ancestry in reticulate lineages. However, they may underestimate signals of lineage-specific or adaptive introgression. As such, this framework is most effective when interpreted as complementary to analyses of more rapidly evolving genomic regions. This distinction is particularly relevant for the study of specialized metabolism, which is often governed by fast-evolving genes and gene clusters. Phylogenetic inference based solely on such loci can be confounded by paralogy and reticulation, especially in polyploid taxa, reinforcing the need to integrate both conserved and rapidly evolving genomic signals.

Within *Chenopodium*, substantial variation in specialized metabolites has been documented, particularly in *C. quinoa*, where flavonoid and saponin profiles vary across genotypes and environmental conditions (Huan et al. 2022). Interpreting the evolutionary origins of such variation, however, requires a robust understanding of the underlying species history. The shared introgression signals identified here across conserved loci indicate that both tetraploid and hexaploid *C. album* cytotypes derive from a common reticulate background. This provides an important evolutionary context for future analyses of trait-associated genes.

More broadly, these results underscore the importance of integrating phylogenomic approaches based on conserved loci with analyses of rapidly evolving, trait-associated genes. While conserved markers help resolve the backbone of evolutionary history, genes involved in specialized metabolism capture more recent and functionally relevant diversification. Combining these complementary perspectives will be essential for understanding how hybridization and polyploidization contribute to the emergence and regulation of ecologically and agriculturally important traits in *Chenopodium*.

## Conclusion

The evolutionary history of the *C. album* complex is best understood within a reticulate framework shaped by hybridization, polyploidization, and incomplete lineage sorting. By integrating complementary phylogenomic approaches, this study shows that genome-wide discordance within the complex cannot be adequately explained by a strictly bifurcating model alone. Tetraploid and hexaploid *C. album* share strong signals of common reticulate ancestry, consistent with contributions from ancient and potentially unsampled progenitor lineages.

These findings provide a clearer evolutionary context for taxonomy and future analyses related to trait evolution and crop-oriented research in *Chenopodium*. More broadly, this study establishes a roadmap for linking evolutionary history with metabolomics, functional genomics, and physiology to understand how hybridization and genome duplication shape ecologically and agriculturally important traits, including defense, resilience, and edibility.

## Supplementary information

A list of supplementary figures and tables is provided here:

- Supplementary Figures S1–S14

**– Figure S1.** Geographical origin of genome assemblies used in the phylogenomic analyses.
**– Figure S2.** Contig-level assembly metrics across all genomes, grouped by genus.
**– Figure S3.** Scaffold-level assembly metrics across all genomes, grouped by genus.
**– Figure S4.** Reconciliation-based species tree inferred with SpeciesRax from 2,298 BUSCO gene family trees (including single-copy and multicopy genes) across 27 taxa.
**– Figure S5.** Top 20 donor species or internal nodes inferred by SpeciesRax to contribute gene transfers.
**– Figure S6.** Species tree inferred using ASTRAL-Pro3 from 2,298 BUSCO gene family trees.
**– Figure S7.** Coalescent-based species tree of *Chenopodium* inferred using ASTRAL-Pro3 from 2,298 BUSCO gene family trees.
**– Figure S8.** Coalescent-based species tree inferred using ASTRAL-IV from 2,298 decomposed BUSCO gene trees generated with DISCO.
**– Figure S9**. Coalescent-based species tree of *Chenopodium* inferred using ASTRAL-IV from 2,298 decomposed BUSCO gene trees generated with DISCO.
**– Figure S10.** Coalescent-based species tree of *Chenopodium* inferred using ASTRAL-IV from 2,298 decomposed BUSCO gene trees generated with DISCO.
**– Figure S11.** Concatenation-based species tree inferred using IQ-TREE2 from 2,298 decomposed BUSCO gene alignments.
**– Figure S12.** Comparison of phylogenetic network inference using PhyloNet and SNaQ under increasing numbers of reticulations.
**– Figure S13.** Global distribution of *Chenopodium* genomes contributing to the *C. album* reference assembly dcCheAlbu1.1.
**– Figure S14.** Global distribution of *Chenopodium* genomes contributing to the *C. album* reference assembly Ca6-1.
- Supplementary Tables S1–S10

**– Table S1.** Taxonomic identities and assembly data for all sampled taxa.
**– Table S2.** Assembly statistics for all sampled genomes.
**– Table S3.** BUSCO completeness metrics for all assemblies.
**– Table S4.** Plant species represented in the BUSCO eudicot reference dataset.
**– Table S5.** PhyloNet model comparison across reticulation numbers.
**– Table S6.** Comparison of total log probabilities across PhyloNet models.
**– Table S7.** SNaQ model comparison across reticulation numbers.
**– Table S8.** BUSCO loci identified as introgressed in both *C. album* cytotypes.
**– Table S9.** BUSCO loci identified as introgressed only in *C. album* Ca6-1 4x.
**– Table S10.** BUSCO loci identified as introgressed only in *C. album* dcCheAlbu1.1 6x.

## Author contributions

K.E. conceived the study, performed the analyses, and wrote the first draft of the manuscript. J.S. contributed to conceptualization, assisted with analytical design and interpretation, and contributed to writing the manuscript. P.D.C. contributed to conceptualization, guided the analytical framework and manuscript structure, and contributed to writing the manuscript. All authors contributed to discussion of the results and approved the final version of the manuscript.

## AI assistance acknowledgement

AI tools were used only for language editing, clarity, and organization of the manuscript text. All scientific content, analyses, and interpretations were developed and verified by the authors.

## Data availability

Genome assemblies used in this study were downloaded from the NCBI Genome Database. Accession numbers are provided in Supplementary Table S1. The tetraploid *C. album* Ca6-1 genome assembly was deposited in NCBI under BioProject PRJNA137704 (Vats et al. 2026). Scripts used for the different analyses are publicly available at the following GitHub repositories: ASTRAL-Pro3 script and Phylogenomics Pipeline script.

