## Supplementary data for "Phylogenomics reveals reticulate evolution in the *Chenopodium album* complex"

**Supplementary Table S1. Taxonomic identities and assembly data for all sampled taxa.**

| Family | Genus | Assembly Accession | Assembly Name | Organism Name | Country | Ploidy |
|---|---|---|---|---|---|---|
| Amaranthaceae | *Beta* | GCF_026745355.1 | EL10.2 | *Beta vulgaris subsp. vulgaris* | USA | 2x |
| Amaranthaceae | *Beta* | GCA_037177695.1 | USDA_Bvulg_FC309_v1.1.0 | *Beta vulgaris subsp. vulgaris* | USA | 2x |
| Amaranthaceae | *Beta* | GCA_947646795.1 | BlomONT_v1.0 | *Beta lomatogona* | Turkey | 2x |
| Amaranthaceae | *Beta* | GCA_947648045.1 | BcorONT_v1.0 | *Beta corolliflora* | Turkey | 4x |
| Amaranthaceae | *Beta* | GCA_947644855.1 | Bmrh_v1.0 | *Beta macrorhiza* | Turkey | 2x |
| Amaranthaceae | *Spinacia* | GCF_020520425.1 | BTI_SOV_V1 | *Spinacia oleracea* | Uknown | 2x |
| Amaranthaceae | *Spinacia* | GCA_019972175.1 | SOL_r1.1 | *Spinacia oleracea* | Japan | 2x |
| Amaranthaceae | *Chenopodium* | Ca6-1[1] | Ca6-1 | *Chenopodium album* | Denmark | 4x |
| Amaranthaceae | *Chenopodium* | GCA_948465745.1 | dcCheAlbu1.1 | *Chenopodium album* | UK | 6x |
| Amaranthaceae | *Chenopodium* | GCA_046718445.1 | ASM4671844v1 | *Chenopodium ficifolium* | USA | 2x |
| Amaranthaceae | *Chenopodium* | GCA_046866025.1 | V1 | *Chenopodium ficifolium* | USA | 2x |
| Amaranthaceae | *Chenopodium* | GCA_046718525.1 | ASM4671852v1 | *Chenopodium opulifolium* | Spain | 6x |
| Amaranthaceae | *Chenopodium* | GCA_046718595.1 | ASM4671859v1 | *Chenopodium sosnowskyi* | Iran | 4x |
| Amaranthaceae | *Chenopodium* | GCA_046718605.1 | ASM4671860v1 | *Chenopodium strictum* | Czech Republic | 4x |
| Amaranthaceae | *Chenopodium* | GCA_046718705.1 | ASM4671870v1 | *Chenopodium acuminatum* | Russia | 2x |
| Amaranthaceae | *Chenopodium* | GCA_024500155.1 | ASM2450015v1 | *Chenopodium formosanum* | Taiwan | 6x |

*Continued on next page*

[1] BioProject PRJNA137704

**Supplementary Table S1. Taxonomic identities and assembly data for all sampled taxa.**

| Family | Genus | Assembly Accession | Assembly Name | Organism Name | Country | Ploidy |
|---|---|---|---|---|---|---|
| Amaranthaceae | *Chenopodium* | GCA_046718575.1 | ASM4671857v1 | *Chenopodium vulvaria* | Hungary | 2x |
| Amaranthaceae | *Chenopodium* | GCA_046718655.1 | ASM4671865v1 | *Chenopodium pamiricum* | Russia | 2x |
| Amaranthaceae | *Chenopodium* | GCA_034703905.1 | C_berlandieri_1.0 | *Chenopodium berlandieri* | Mexico | 4x |
| Amaranthaceae | *Chenopodium* | GCA_040571465.1 | ASM4057146v1 | *Chenopodium quinoa* | Bolivia | 4x |
| Amaranthaceae | *Chenopodium* | GCA_001687005.1 | ASM168700v1 | *Chenopodium pallidicaule* | USA | 2x |
| Amaranthaceae | *Chenopodium* | GCA_001687025.1 | ASM168702v1 | *Chenopodium suecicum* | USA | 2x |
| Amaranthaceae | *Amaranthus* | GCA_019425755.1 | ASM1942575v1 | *Amaranthus cruentus* | Tanzania | 2x |
| Amaranthaceae | *Amaranthus* | GCA_000753965.2 | IBAB_Ahyp_2.0 | *Amaranthus hypochondriacus* | India | 2x |
| Amaranthaceae | *Amaranthus* | GCA_019776075.1 | ASM1977607v1 | *Amaranthus palmeri* | USA | 2x |
| Amaranthaceae | *Amaranthus* | GCF_026212465.1 | ASM2621246v1 | *Amaranthus tricolor* | China | 2x |
| Caryphyllaceae | *Dianthus* | GCA_000512335.1 | DCA_r1.0 | *Dianthus caryophyllus* | Mediterranean | 2x |

**Supplementary Table S2. Assembly statistics for all sampled genomes.**

| Assembly Accession | No. Scaffolds | Scaffold N50 | Scaffold L50 | No. Contigs | Contig N50 | Contig L50 | GC (%) | Assembly Level |
|---|---|---|---|---|---|---|---|---|
| GCF_026745355.1 | 18 | 62 Mb | 5 | 3098 | 1.3 Mb | 119 | 36 | Chromosome |
| GCA_037177695.1 | 260 | 67.8 Mb | 5 | 293 | 31.5 Mb | 8 | 36 | Chromosome |
| GCA_947646795.1 | – | – | – | 1530 | 1.7 Mb | 141 | 365 | Contig |
| GCA_947648045.1 | – | – | – | 4355 | 720.7 kb | 583 | 37 | Contig |
| GCA_947644855.1 | 218216 | 7.1 kb | 23482 | 222966 | 6.7 kb | 27985 | 365 | Scaffold |
| GCF_020520425.1 | 244 | 151.5 Mb | 3 | 307 | 23.8 Mb | 12 | 38 | Chromosome |
| GCA_019972175.1 | 287 | 11.3 Mb | 25 | 17256 | 182.5 kb | 1475 | 38 | Scaffold |
| Ca6-1 | – | – | – | 1261 | 50.42 Mb | 14 | 36 | Contig |
| GCA_948465745.1 | 156 | 60.3 Mb | 11 | 223 | 31.5 Mb | 17 | 36 | Chromosome |
| GCA_046718445.1 | 324 | 79.9 Mb | 5 | 326 | 79.9 Mb | 5 | 37 | Chromosome |
| GCA_046866025.1 | 777 | 79.9 Mb | 5 | 779 | 79.9 Mb | 5 | 37 | Chromosome |
| GCA_046718525.1 | 309 | 70.8 Mb | 12 | 310 | 66.3 Mb | 12 | 365 | Chromosome |
| GCA_046718595.1 | 276 | 53.4 Mb | 8 | 286 | 43 Mb | 10 | 36 | Chromosome |
| GCA_046718605.1 | 252 | 51.6 Mb | 8 | 259 | 38.6 Mb | 8 | 355 | Chromosome |
| GCA_046718705.1 | 180 | 45.3 Mb | 5 | 190 | 34 Mb | 6 | 355 | Chromosome |
| GCA_024500155.1 | 798 | 61.1 Mb | 11 | 846 | 32.6 Mb | 18 | 36 | Chromosome |
| GCA_046718575.1 | 379 | 41.3 Mb | 5 | 396 | 22.2 Mb | 7 | 35 | Chromosome |
| GCA_046718655.1 | 949 | 51.6 Mb | 5 | 995 | 16.7 Mb | 9 | 37 | Chromosome |
| GCA_034703905.1 | 336 | 70.1 Mb | 9 | 1191 | 4.9 Mb | 76 | 375 | Chromosome |
| GCA_040571465.1 | 18 | 71.1 Mb | 9 | 161 | 29.4 Mb | 16 | 37 | Chromosome |

*Continued on next page*

| Assembly Accession | No. Scaffolds | Scaffold N50 | Scaffold L50 | No. Contigs | Contig N50 | Contig L50 | GC (%) | Assembly Level |
|---|---|---|---|---|---|---|---|---|
| GCA_001687005.1 | 3013 | 356.8 kb | 243 | 9244 | 78.5 kb | 1174 | 365 | Scaffold |
| GCA_001687025.1 | 11198 | 105.4 kb | 1285 | 38616 | 27.4 kb | 5069 | 36 | Scaffold |
| GCA_019425755.1 | 625 | 21.7 Mb | 8 | 1608 | 898.7 kb | 86 | 33 | Chromosome |
| GCA_000753965.2 | 1567 | 22.6 Mb | 8 | 8151 | 82.8 kb | 1340 | 33 | Chromosome |
| GCA_019776075.1 | – | – | – | 1664 | 601.1 kb | 145 | 335 | Contig |
| GCF_026212465.1 | 48 | 31.7 Mb | 8 | 2544 | 905.9 kb | 99 | 32 | Chromosome |
| GCA_000512335.1 | 45088 | 60.7 kb | 2075 | 89083 | 16.7 kb | 7147 | 365 | Scaffold |

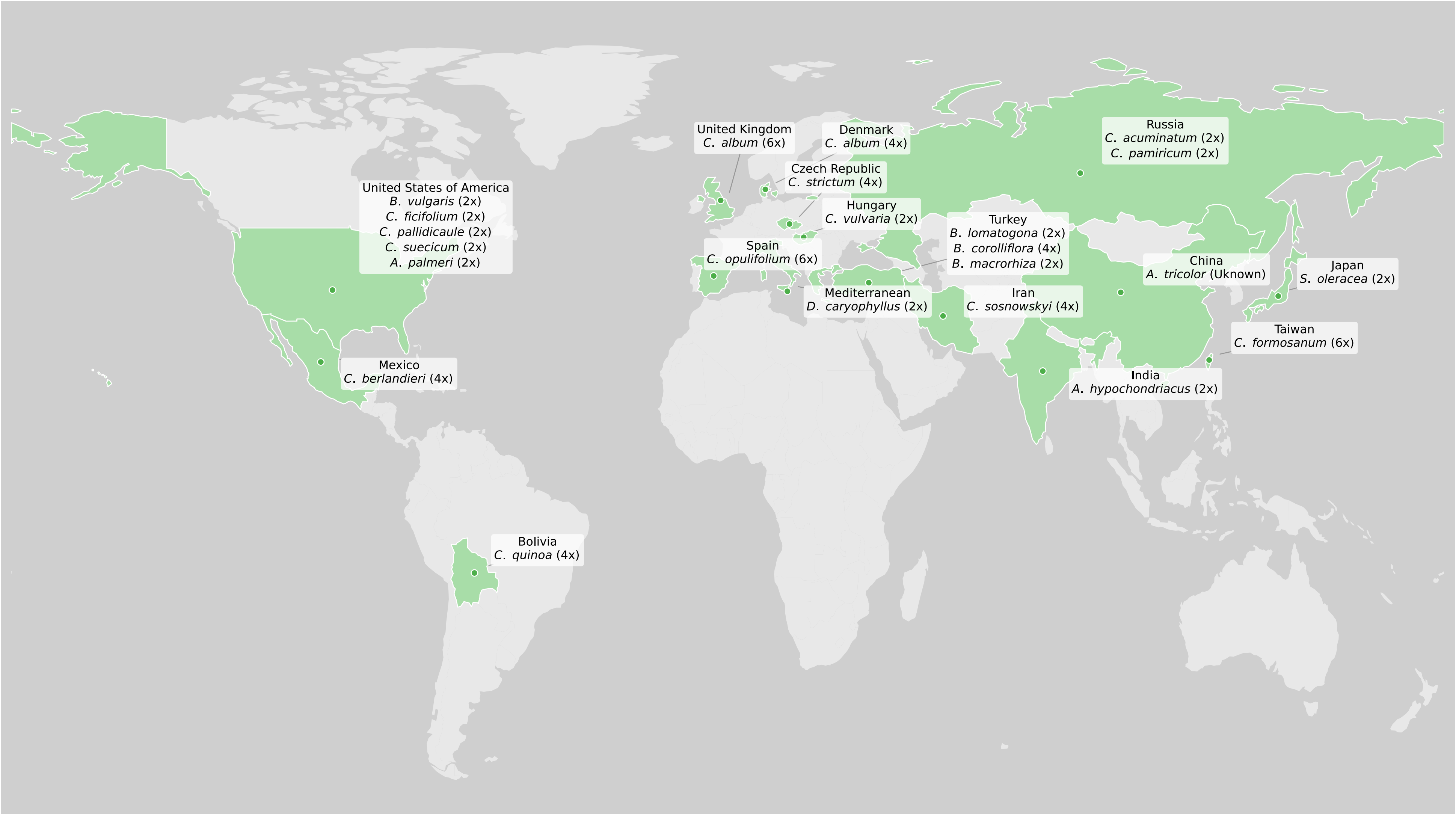


**Supplementary Figure S1. Geographical origin of genome assemblies used in the phylogenomic analyses.** World map showing the country of origin for each genome assembly included in this study. Labels indicate species identity and ploidy level. The dataset includes representatives of *Chenopodium*, *Spinacia*, *Beta*, *Amaranthus*, and *Dianthus*. Tetraploid and hexaploid cytotypes of *Chenopodium album* are represented by genomes from Denmark (4x) and the United Kingdom (6x), respectively.

**Supplementary Table S3. BUSCO completeness metrics for all assemblies.** C: Complete, S: Single-copy, D: Duplicated, F: Fragmented, M: Missing.

| Assembly Accession | Assembly Name | Organism Name | Ploidy | C | S | D | F | M |
|---|---|---|---|---|---|---|---|---|
| Ca6-1 | Ca6-1 | *Chenopodium album* | tetraploid | 2285 | 47 | 2238 | 5 | 36 |
| GCF_026745355.1 | EL10.2 | *Beta vulgaris* | diploid | 2233 | 2193 | 40 | 14 | 79 |
| GCA_037177695.1 | USDA_Bvulg_FC309_v1.1.0 | *Beta vulgaris* | diploid | 2268 | 2223 | 45 | 10 | 48 |
| GCA_947646795.1 | BlomONT_v1.0 | *Beta lomatogona* | diploid | 2184 | 1272 | 912 | 16 | 126 |
| GCA_947648045.1 | BcorONT_v1.0 | *Beta corolliflora* | tetraploid | 2267 | 329 | 1938 | 12 | 47 |
| GCA_947644855.1 | Bmrh_v1.0 | *Beta macrorhiza* | diploid | 1666 | 1559 | 107 | 340 | 320 |
| GCF_020520425.1 | BTI_SOV_V1 | *Spinacia oleracea* | diploid | 2251 | 2178 | 73 | 13 | 62 |
| GCA_019972175.1 | SOL_r1.1 | *Spinacia oleracea* | diploid | 2239 | 2129 | 110 | 17 | 70 |
| GCA_948465745.1 | dcCheAlbu1.1 | *Chenopodium album* | hexaploid | 2285 | 48 | 2237 | 5 | 36 |
| GCA_046718445.1 | ASM4671844v1 | *Chenopodium ficifolium* | diploid | 2259 | 2210 | 49 | 12 | 55 |
| GCA_046866025.1 | V1 | *Chenopodium ficifolium* | diploid | 2259 | 2210 | 49 | 12 | 55 |
| GCA_046718525.1 | ASM4671852v1 | *Chenopodium opulifolium* | hexaploid | 2280 | 42 | 2238 | 5 | 41 |
| GCA_046718595.1 | ASM4671859v1 | *Chenopodium sosnowskyi* | tetraploid | 2279 | 163 | 2116 | 5 | 42 |
| GCA_046718605.1 | ASM4671860v1 | *Chenopodium strictum* | tetraploid | 2279 | 142 | 2137 | 5 | 42 |
| GCA_046718705.1 | ASM4671870v1 | *Chenopodium acuminatum* | diploid | 2262 | 2200 | 62 | 9 | 55 |
| GCA_024500155.1 | ASM2450015v1 | *Chenopodium formosanum* | hexaploid | 2287 | 52 | 2235 | 6 | 33 |
| GCA_046718575.1 | ASM4671857v1 | *Chenopodium vulvaria* | diploid | 2268 | 2225 | 43 | 9 | 49 |
| GCA_046718655.1 | ASM4671865v1 | *Chenopodium pamiricum* | diploid | 2263 | 2174 | 89 | 9 | 54 |
| GCA_034703905.1 | C.berlandieri_1.0 | *Chenopodium berlandieri* | tetraploid | 2280 | 179 | 2102 | 4 | 42 |
| GCA_001687005.1 | ASM168700v1 | *Chenopodium pallidicaule* | diploid | 2256 | 2213 | 43 | 13 | 57 |

*Continued on next page*

| Assembly Accession | Assembly Name | Organism Name | Ploidy | C | S | D | F | M |
|---|---|---|---|---|---|---|---|---|
| GCA_001687025.1 | ASM168702v1 | *Chenopodium suecicum* | diploid | 2251 | 2204 | 47 | 18 | 57 |
| GCA_040571465.1 | ASM4057146v1 | *Chenopodium quinoa* | tetraploid | 2281 | 218 | 2063 | 4 | 41 |
| GCA_019425755.1 | ASM1942575v1 | *Amaranthus cruentus* | diploid | 2155 | 2060 | 95 | 16 | 155 |
| GCA_000753965.2 | IBAB_Ahyp_2.0 | *Amaranthus hypochondriacus* | diploid | 2018 | 1799 | 219 | 90 | 218 |
| GCA_019776075.1 | ASM1977607v1 | *Amaranthus palmeri* | diploid | 2228 | 1930 | 298 | 16 | 82 |
| GCF_026212465.1 | ASM2621246v1 | *Amaranthus tricolor* | diploid | 2229 | 2116 | 113 | 12 | 85 |
| GCA_000512335.1 | DCA_r1.0 | *Dianthus caryophyllus* | diploid | 2180 | 1549 | 631 | 19 | 127 |

**Supplementary Table S4. Plant species represented in the BUSCO eudicot reference dataset.** Conserved ortholog groups from these taxa were used to assess genome completeness using profile hidden Markov model searches implemented in HMMER.

| NCBI Taxonomy ID | Species |
|---:|---|
| 200316 | *Nicotiana obtusifolia* |
| 3711 | *Brassica rapa* |
| 4155 | *Erythranthe guttata* |
| 3988 | *Ricinus communis* |
| 3641 | *Theobroma cacao* |
| 3775 | *Cephalotus follicularis* |
| 3555 | *Beta vulgaris subsp. vulgaris* |
| 3885 | *Phaseolus vulgaris* |
| 3983 | *Manihot esculenta* |
| 3702 | *Arabidopsis thaliana* |
| 49451 | *Nicotiana attenuata* |
| 33119 | *Petunia axillaris* |
| 4236 | *Lactuca sativa* |
| 57577 | *Trifolium pratense* |
| 81972 | *Arabidopsis lyrata subsp. lyrata* |
| 51240 | *Juglans regia* |
| 29760 | *Vitis vinifera* |
| 4072 | *Capsicum annuum* |
| 29730 | *Gossypium raimondii* |
| 3914 | *Vigna angularis* |
| 71139 | *Eucalyptus grandis* |
| 3827 | *Cicer arietinum* |
| 4113 | *Solanum tuberosum* |
| 2711 | *Citrus sinensis* |
| 3656 | *Cucumis melo* |
| 3760 | *Prunus persica* |
| 72664 | *Eutrema salsugineum* |
| 81985 | *Capsella rubella* |
| 4081 | *Solanum lycopersicum* |
| 4039 | *Daucus carota* |
| 101020 | *Fragaria vesca subsp. vesca* |

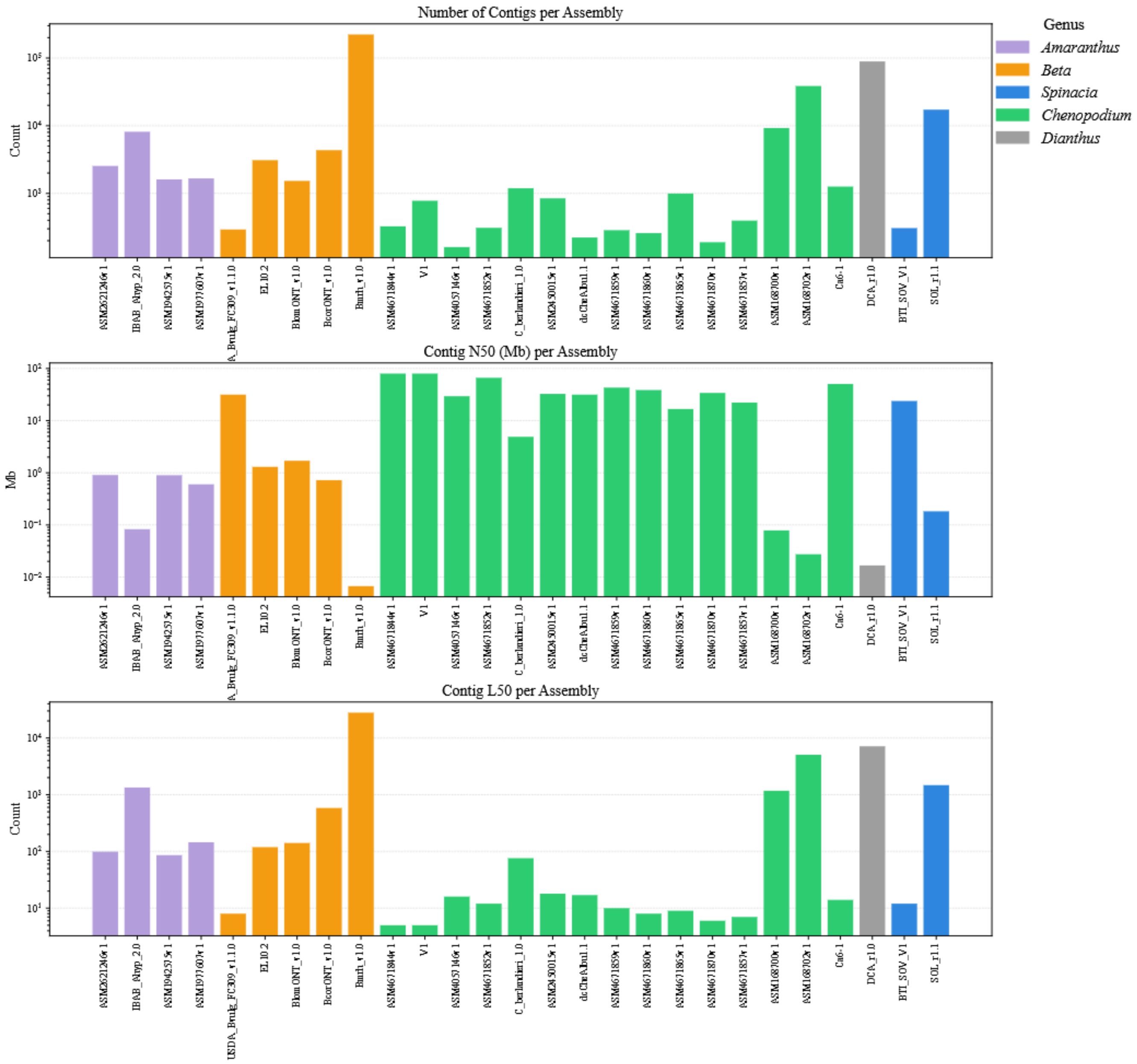


**Supplementary Figure S2. Contig-level assembly metrics across all genomes, grouped by genus.** The top panel shows the number of contigs per assembly, the middle panel shows contig N50 (Mb), and the bottom panel shows contig L50. Logarithmic scaling is applied where necessary to accommodate the wide range of values. Colours represent genus-level classification. As expected, contig counts are generally higher than scaffold counts, reflecting fragmentation prior to scaffolding. Contig N50 values show substantial variability across assemblies, with high-quality genomes characterized by low contig counts, high N50 values, and low L50 values. In contrast, fragmented assemblies exhibit high contig counts and low N50. Assemblies from *Chenopodium* again display relatively consistent metrics, while other genera show broader variation.

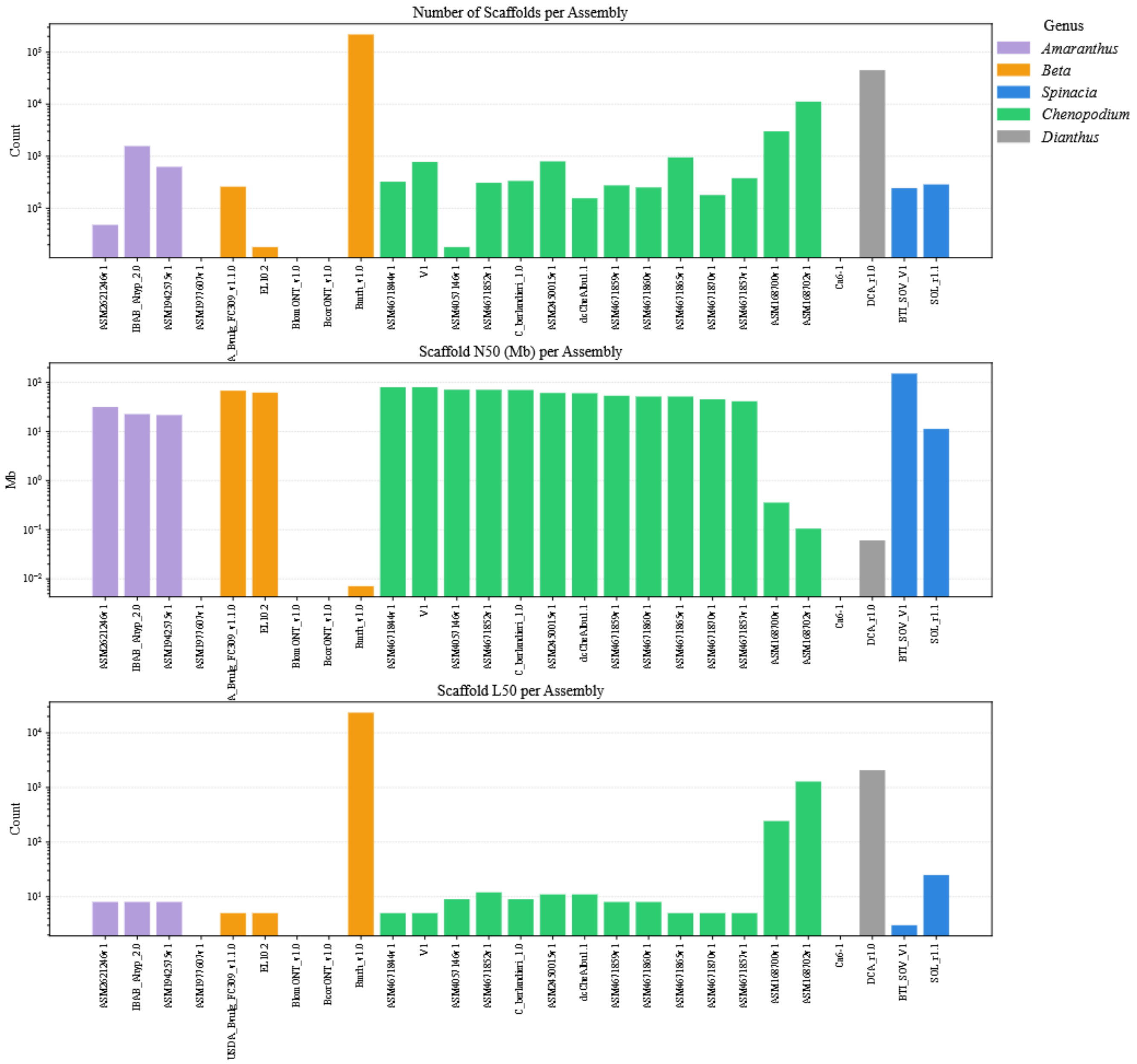


**Supplementary Figure S3. Scaffold-level assembly metrics across all genomes, grouped by genus.** The top panel shows the number of scaffolds per assembly, the middle panel shows scaffold N50 (Mb), and the bottom panel shows scaffold L50. Values are displayed on a logarithmic scale where appropriate to highlight variation across assemblies. Colours denote genus-level classification. Substantial variation in assembly contiguity is observed across taxa. Assemblies with fewer scaffolds generally exhibit higher N50 values and lower L50, indicating high contiguity. In contrast, highly fragmented assemblies show low N50 and high L50 values. Assemblies within the genus *Chenopodium* have high consistent assembly quality, whereas other genera (e.g., *Beta* and *Dianthus*) display lower contiguity. Assemblies lacking bars correspond to genomes that were not scaffolded, and therefore do not have scaffold-level metrics.

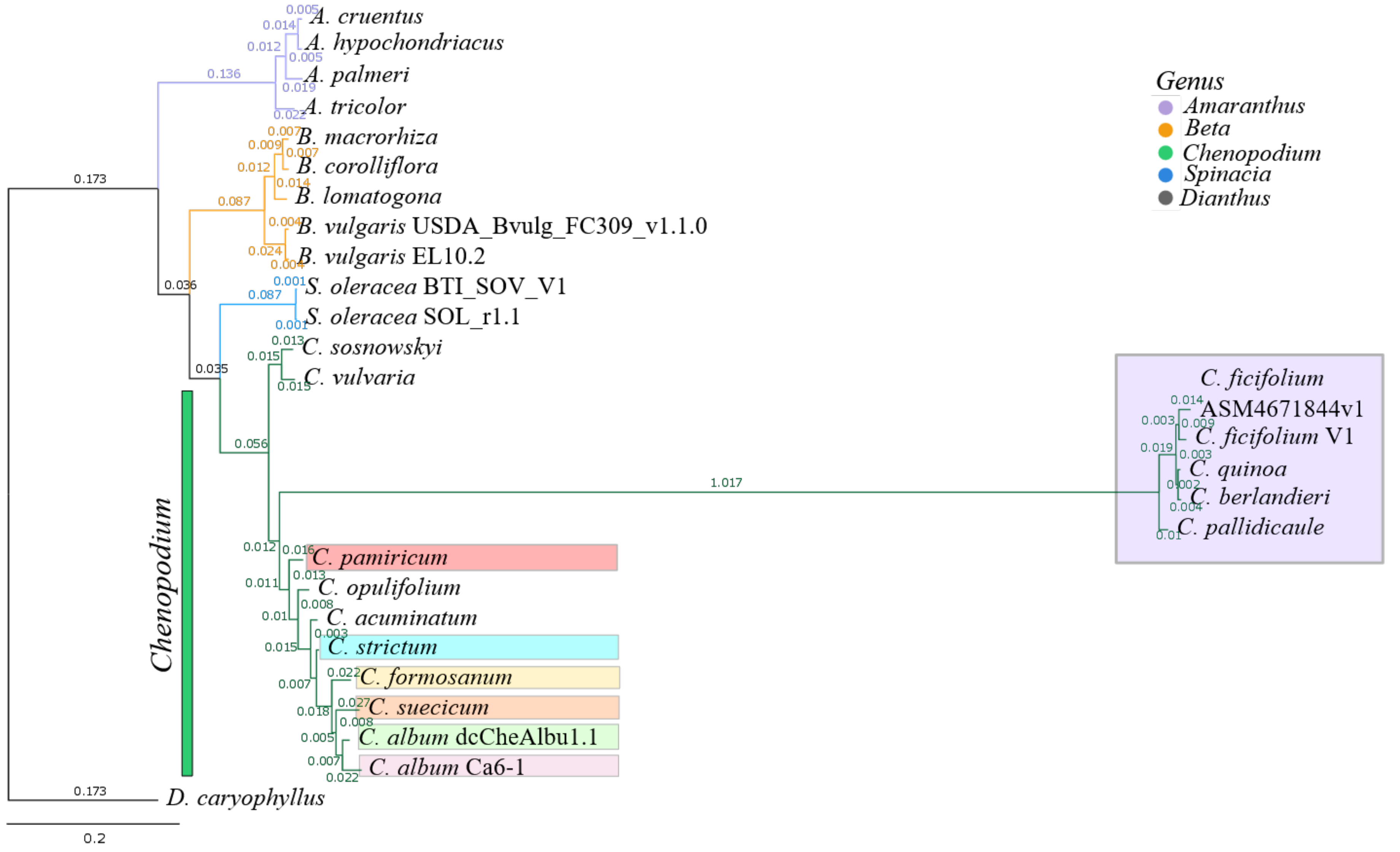


**Supplementary Figure S4. Reconciliation-based species tree inferred with SpeciesRax from 2,298 BUSCO gene family trees (including single-copy and multi-copy genes) across 27 taxa.** SpeciesRax estimates a rooted species tree under the UndatedDTL model by jointly optimizing the likelihood of gene family evolution with respect to gene duplication, loss, and gene transfer events. Branch lengths represent the expected number of substitutions per site. Node support values correspond to speciation-driven quartets (SQs), reported as QPIC/EQPIC scores, where values close to 1 indicate strong support for the inferred bipartition and negative values indicate support for alternative topologies.

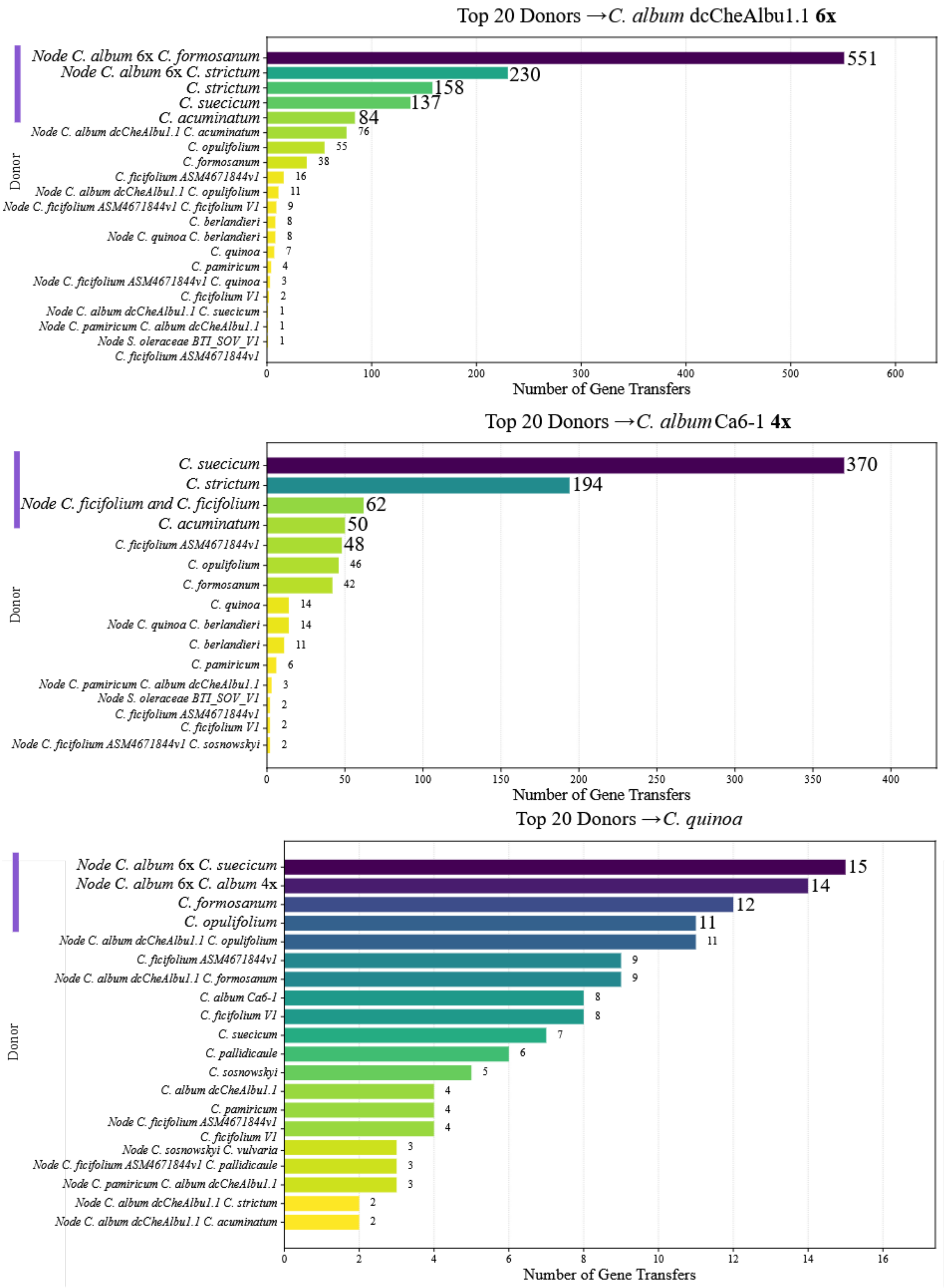


**Supplementary Figure S5. Top 20 donor species or internal nodes inferred by SpeciesRax to contribute gene transfers.** Bar plots show the number of inferred transfer events to *C. album* dcCheAlbu1.1 (6x; top), *C. album* Ca6-1 (4x; middle), and *C. quinoa* (bottom). Bars corresponding to the highest contributing donors in each panel are highlighted with a thick purple line. Donors correspond either to extant taxa or to internal nodes of the species tree. "Node" represents internal nodes defined by the most recent common ancestor of the listed taxa.

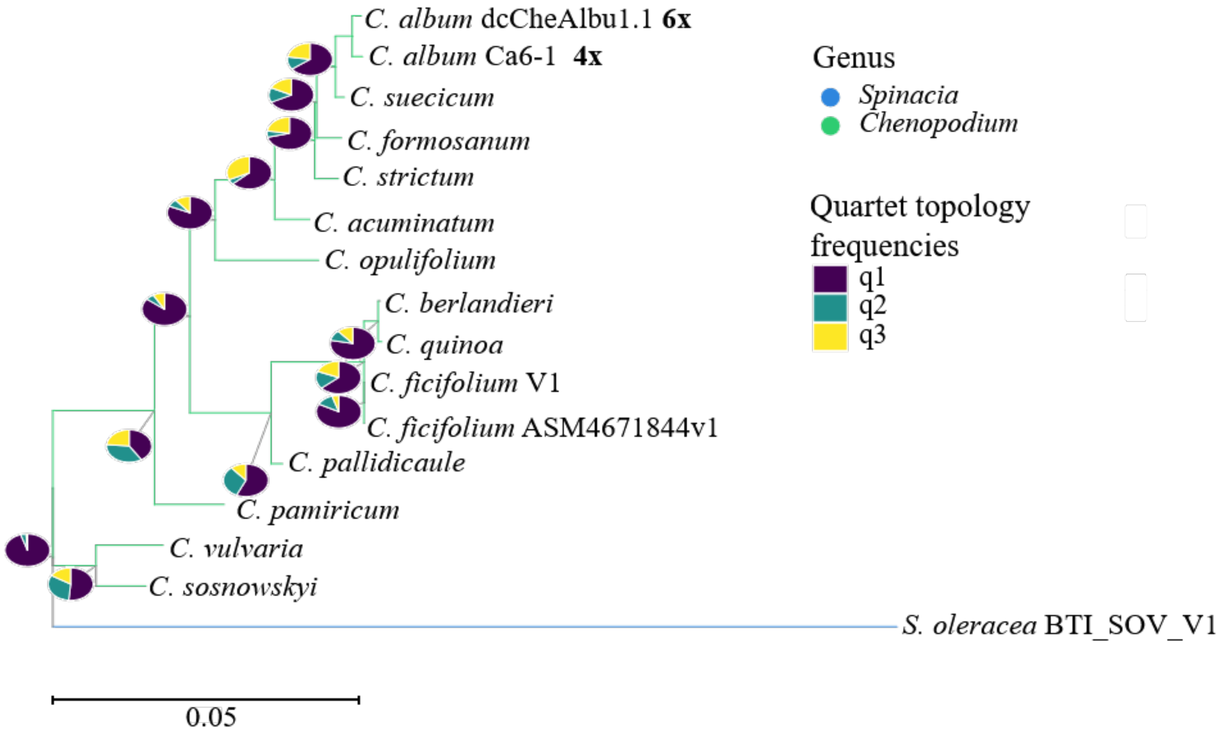


**Supplementary Figure S6. Species tree inferred using ASTRAL-Pro3 from 2,298 BUSCO gene family trees.** The phylogeny includes 16 taxa, with *S. oleracea* used as the outgroup. Branch lengths are shown in coalescent units. Node support is represented by pie charts – the relative frequencies of the three alternative quartet topologies (q1–q3). Most nodes are strongly supported by a dominant quartet topology (q1). However, several internal nodes display more evenly distributed quartet frequencies, indicating localized gene tree discordance. The genus *Chenopodium* forms a well-resolved clade, with closely related species such as *C. album* accessions clustering together. The placement of taxa within this clade is largely consistent, although some nodes exhibit reduced support.

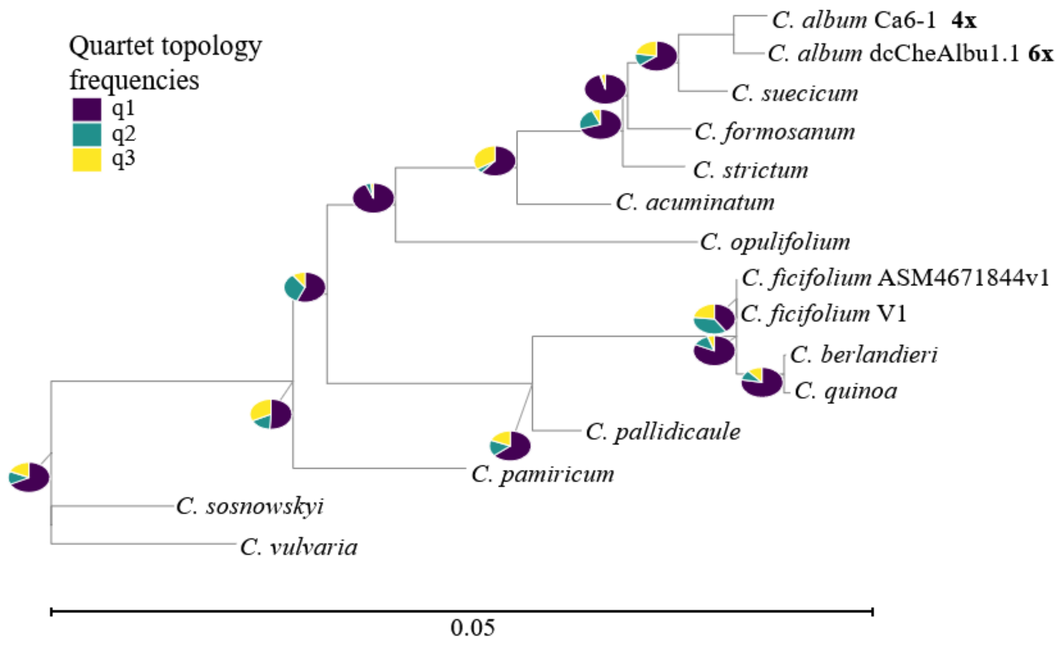


**Supplementary Figure S7. Coalescent-based species tree of *Chenopodium* inferred using ASTRAL-Pro3 from 2,298 BUSCO gene family trees.** The phylogeny includes 15 taxa, with branch lengths shown in coalescent units. The tree resolves major relationships within *Chenopodium*, with closely related taxa clustering together. Most nodes are also supported by a dominant quartet topology. A similar pattern is observed in the genus-level phylogeny that includes *Spinacia* as an outgroup.

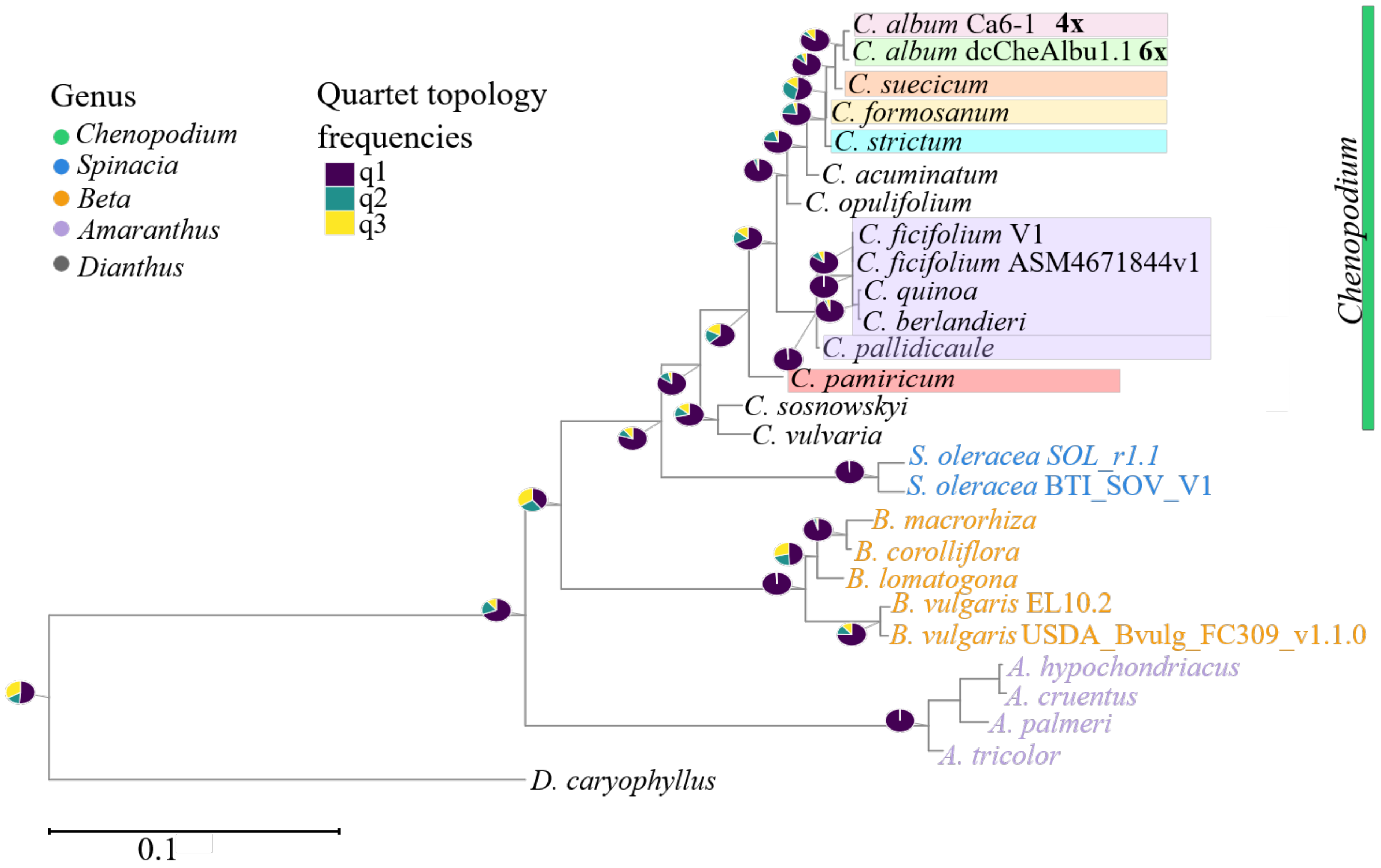


**Supplementary Figure S8. Coalescent-based species tree inferred using ASTRAL-IV from 2,298 decomposed BUSCO gene trees generated with DISCO.** The phylogeny includes 27 taxa, with *Dianthus caryophyllus* used as the outgroup. Branch lengths are shown in coalescent units. Pie charts at internal nodes represent the relative frequencies of the main and two alternative quartet topologies (q1–q3). The tree recovers the same major topology as the species trees inferred using ASTRAL-Pro3. However, several deeper nodes still exhibit mixed quartet frequencies, suggesting that gene tree discordance persists despite the use of decomposed gene trees.

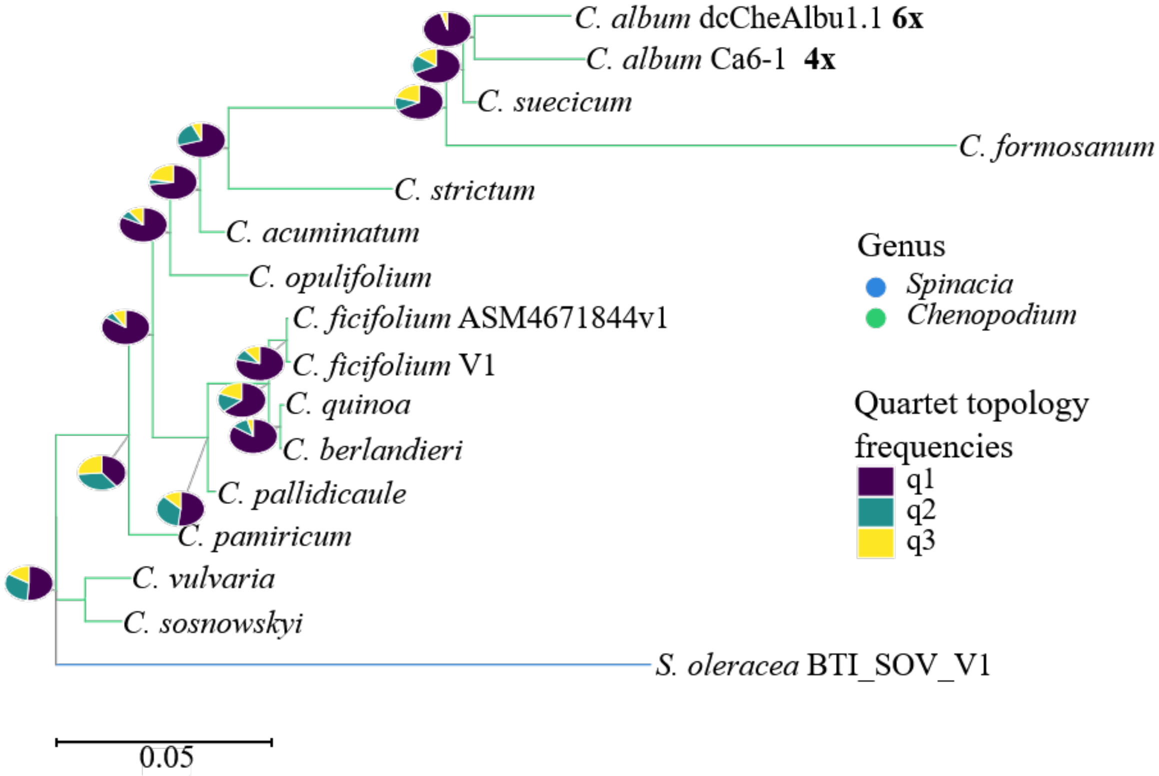


**Supplementary Figure S9. Coalescent-based species tree of *Chenopodium* inferred using ASTRAL-IV from 2,298 decomposed BUSCO gene trees generated with DISCO.** The phylogeny includes 16 taxa, with *S. oleracea* used as the outgroup. Branch lengths are shown in coalescent units. Pie charts at internal nodes represent the relative frequencies of the main and two alternative quartet topologies (q1–q3). The tree recovers the same major relationships within *Chenopodium* as observed in analyses using ASTRAL-Pro3 and ASTRAL-IV with *D. caryophyllus* as outgroup, indicating consistency across methods and datasets. Most nodes are supported by a dominant quartet topology and several internal nodes still exhibit mixed quartet frequencies.

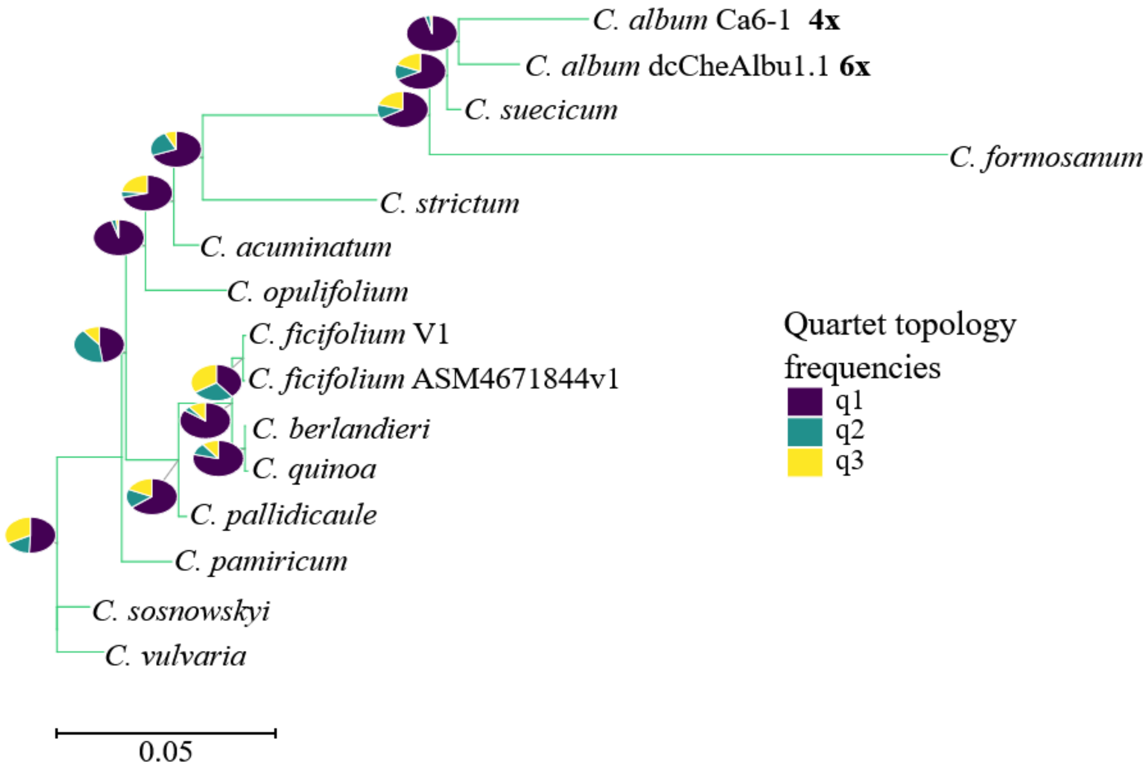


**Supplementary Figure S10. Coalescent-based species tree of *Chenopodium* inferred using ASTRAL-IV from 2,298 decomposed BUSCO gene trees generated with DISCO.** The phylogeny includes 15 *Chenopodium* taxa. The tree resolves relationships within *Chenopodium*, with closely related taxa forming a clade. The overall topology is consistent with previous analyses.

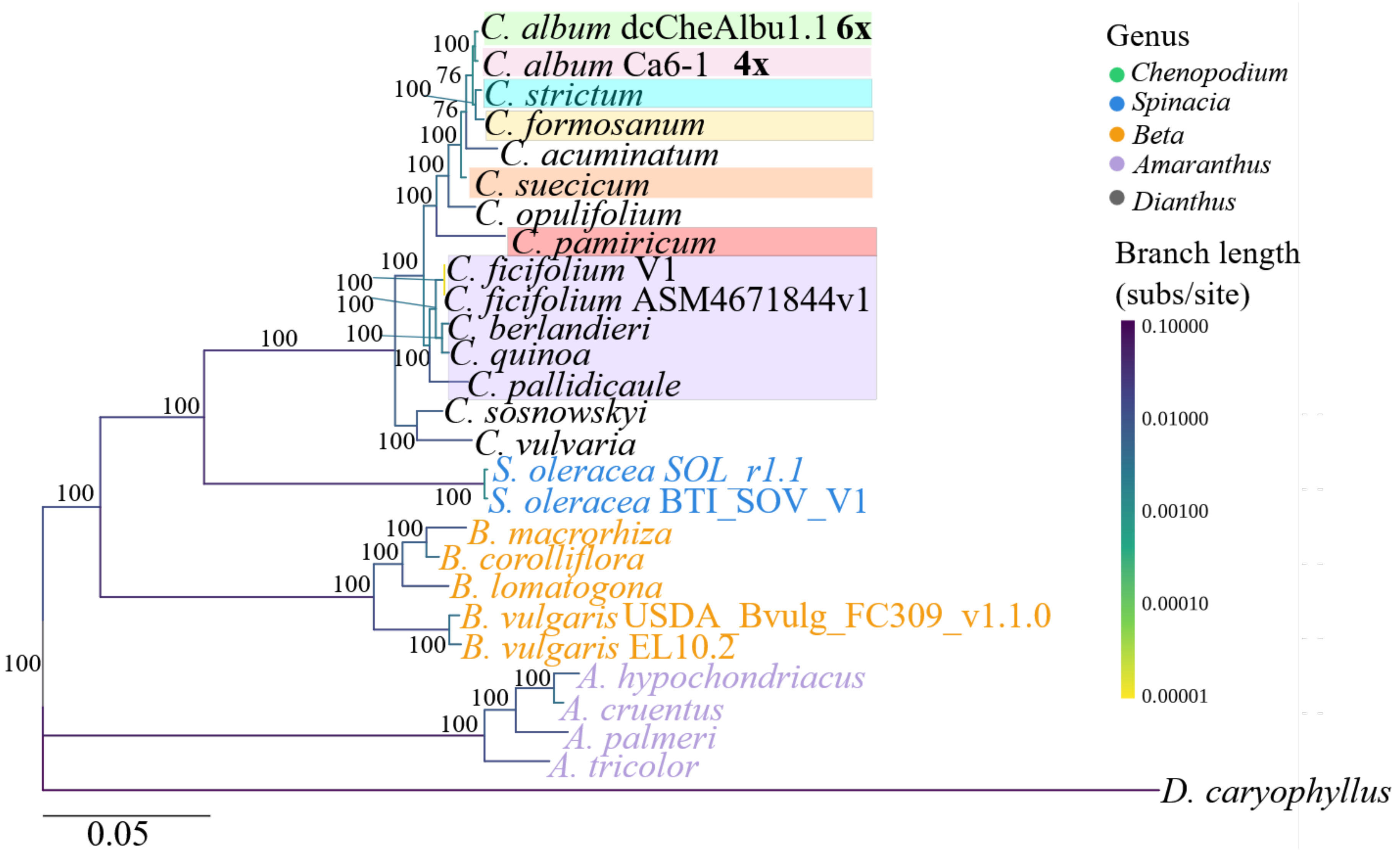


**Supplementary Figure S11. Concatenation-based species tree inferred using IQ-TREE2 from 2,298 decomposed BUSCO gene alignments.** The phylogeny includes 27 taxa, with *D. caryophyllus* used as the outgroup. Branch lengths are expressed in substitutions per site, and node labels indicate bootstrap support values. The tree resolves major relationships across genera, with clear clustering of taxa within *Chenopodium*, *Spinacia*, *Beta*, and *Amaranthus*. The overall topology of *C. album* clade is largely inconsistent with the coalescent-based trees inferred using coalescent-based inference. This approach does not explicitly account for gene tree discordance.

**Supplementary Table S5. PhyloNet model comparison across reticulation numbers.** Phylogenetic network models were inferred using PhyloNet under varying numbers of allowed reticulation events ($h$ = 0–3), with three independent runs performed for each value of $h$ to assess convergence and *S. oleracea* as the outgroup. For each run, pseudolikelihood scores, number of taxa, number of inferred parameters ($k$), number of sites, and model selection criteria (AIC and BIC) are reported. Higher (less negative) pseudolikelihood values indicate better model fit, while AIC and BIC penalize model complexity to enable comparison across models with different numbers of reticulations. Across all runs, models allowing reticulation ($h > 0$) show improved fit compared to strictly tree-like models ($h = 0$). The best-supported model based on AIC and BIC corresponds to $h = 3$. Consistency across replicate runs further indicates stable inference under this parameter setting.

| *h* | *run* | Pseudolikelihood | No. of taxa | k | No.of sites | AIC | BIC |
|---|---|---|---|---|---|---|---|
| 0 | 1 | -242865.9809 | 16 | 15 | 2298 | 485761.9618 | 485848.0587 |
| 0 | 2 | -242865.9809 | 16 | 15 | 2298 | 485761.9618 | 485848.0587 |
| **0** | **3** | **-242865.9809** | **16** | **15** | **2298** | **485761.9618** | **485848.0587** |
| 1 | 1 | -239188.5725 | 16 | 17 | 2298 | 478411.1450 | 478508.7215 |
| 1 | 2 | -239188.5725 | 16 | 17 | 2298 | 478411.1450 | 478508.7215 |
| **1** | **3** | **-239188.5725** | **16** | **17** | **2298** | **478411.1450** | **478508.7215** |
| 2 | 1 | -241349.9059 | 16 | 19 | 2298 | 482737.8118 | 482846.8679 |
| 2 | 2 | -236006.5041 | 16 | 19 | 2298 | 472051.0082 | 472160.0643 |
| **2** | **3** | **-235706.0183** | **16** | **19** | **2298** | **471450.0366** | **471559.0927** |
| 3 | 1 | -235243.5871 | 16 | 21 | 2298 | 470529.1742 | 470649.7099 |
| 3 | 2 | -235243.5871 | 16 | 21 | 2298 | 470529.1742 | 470649.7099 |
| **3** | **3** | **-235243.5871** | **16** | **21** | **2298** | **470529.1742** | **470649.7099** |

**Supplementary Table S6. Comparison of total log probabilities across PhyloNet models.** Total log probabilities are shown for phylogenetic network models inferred under different numbers of reticulation events ($h = 0 - 3$). Increasing the number of allowed reticulations results in improved (less negative) log probability values. The highest likelihood is observed for $h = 3$, although network inferred did not recover the main topology within *C. album* lineage (Supplementary Figure S12).

| *h* | Total log prob | Notes |
|---|---|---|
| 0 | -22680.5381 | shown in the Supplementary Figure S12 |
| 1 | -22197.1267 | shown in the Supplementary Figure S12 |
| 2 | -21468.9852 | shown in the Supplementary Figure S12 |
| 3 | -21684.1686 | shown in the Figure 6 and Supplementary Figure S12 |

**Supplementary Table S7. SNaQ model comparison across reticulation numbers.** Phylogenetic network models were inferred using SNaQ under increasing numbers of allowed reticulation events ($h = 0 - 3$), with multiple independent runs performed for each value of $h$ and *S. oleracea* as the outgroup. For each run, the number of taxa, log-likelihood ($-\log L$), number of parameters ($k$), and model selection criteria (AIC and BIC) are reported. Models allowing reticulation ($h > 0$) show improved fit compared to the tree model ($h = 0$). The best-supported model based on AIC and BIC corresponds to $h = 2$ – a network with two reticulation events provides the optimal balance between fit and complexity. Associated network topologies are shown in Figure 6 and Supplementary Figure S12.

| $h$ | $n_{\text{taxa}}$ | $-\log L$ | $k$ | AIC | BIC | Notes |
|---|---|---|---|---|---|---|
| 0 | 16 | **-25744.4517** | 15 | **-51458.9034** | **-51372.80648** | shown in Supplementary |
| 0 | 16 | -25744.4517 | 15 | -51458.9034 | -51372.80648 | Figure S12 |
| 1 | 16 | -30374.95529 | 17 | -60715.91058 | -60618.33408 | |
| 1 | 16 | **-24857.57341** | 17 | **-49681.14682** | **-49583.57032** | shown in Supplementary |
| 2 | 16 | -20577.3866 | 19 | -41116.77319 | -41007.7171 | Figure S12 |
| 2 | 16 | -22328.28162 | 19 | -44618.56324 | -44509.50715 | |
| 2 | 16 | -25162.51297 | 19 | -50287.02595 | -50177.96985 | |
| 2 | 16 | **-17436.02616** | 19 | **-34834.05231** | **-34724.99622** | shown in Figure 6 |
| 2 | 16 | -19417.07322 | 19 | -38796.14644 | -38687.09034 | and Supplementary |
| 2 | 16 | -26756.79846 | 19 | -53475.59692 | -53366.54082 | Figure S12 |
| 3 | 16 | **-19767.84581** | 21 | **-39493.69163** | **-39373.15595** | shown in Supplementary |
| 3 | 16 | -20028.12351 | 21 | -40014.24702 | -39893.71133 | Figure S12 |
| 3 | 16 | -20244.16137 | 21 | -40446.32275 | -40325.78706 | |
| 3 | 16 | -19088.18863 | 21 | -38134.37726 | -38013.84158 | |
| 3 | 16 | -19859.17376 | 21 | -39676.34752 | -39555.81183 | |
| 3 | 16 | -20017.11947 | 21 | -39992.23894 | -39871.70326 | |

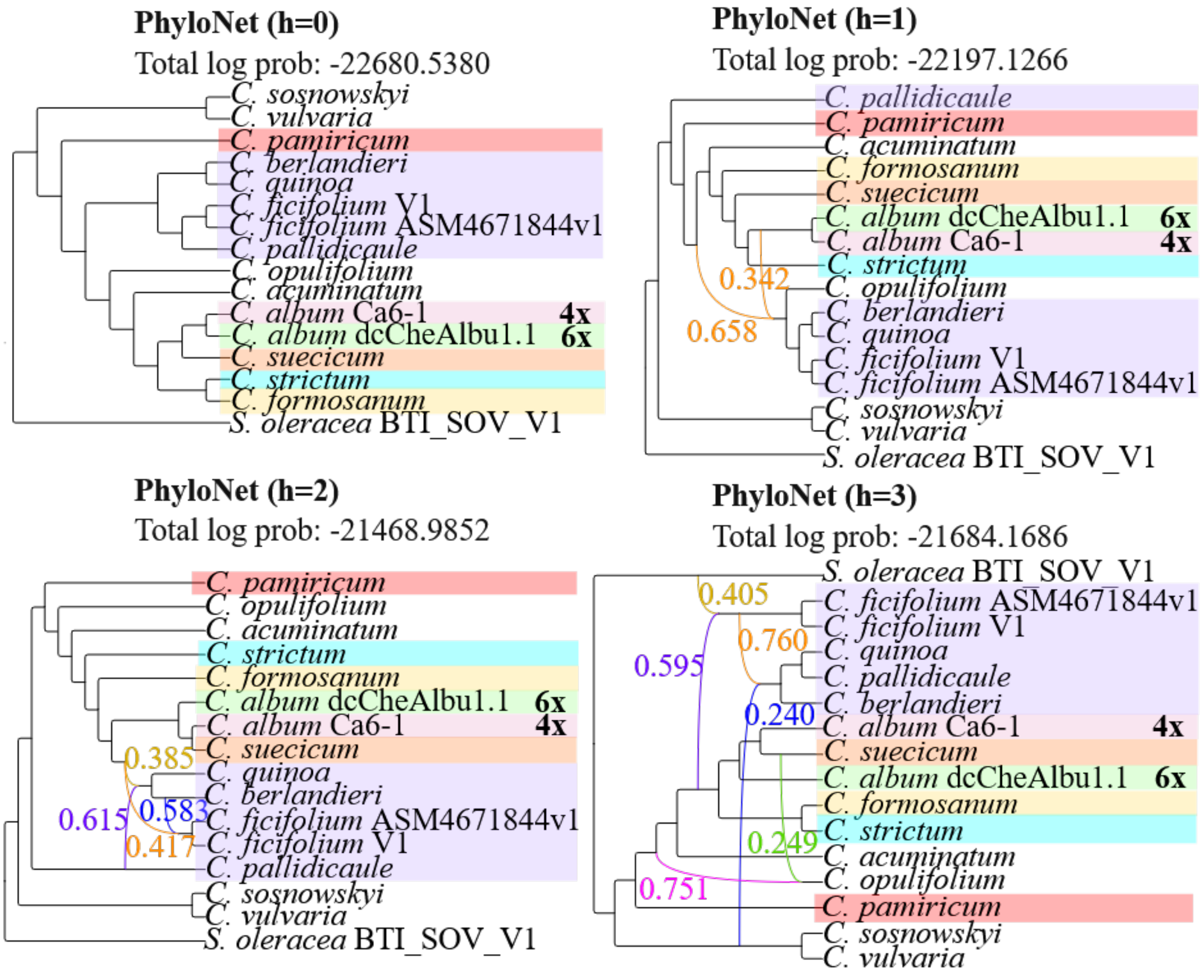


**Supplementary Figure S12. Comparison of phylogenetic network inference using PhyloNet and SNaQ under increasing numbers of reticulations.** (A) PhyloNet networks inferred with reticulation numbers ranging from $h = 0$ to $h = 3$. For each value of $h$, the best-scoring network is shown, with log probabilities indicated. Coloured edges indicate reticulation, where each pair of edges, in complementary colours, sums up to 1.Across all models, major clades are consistently recovered, although the placement of the two *C. album* cytotypes varies with increasing reticulation number.

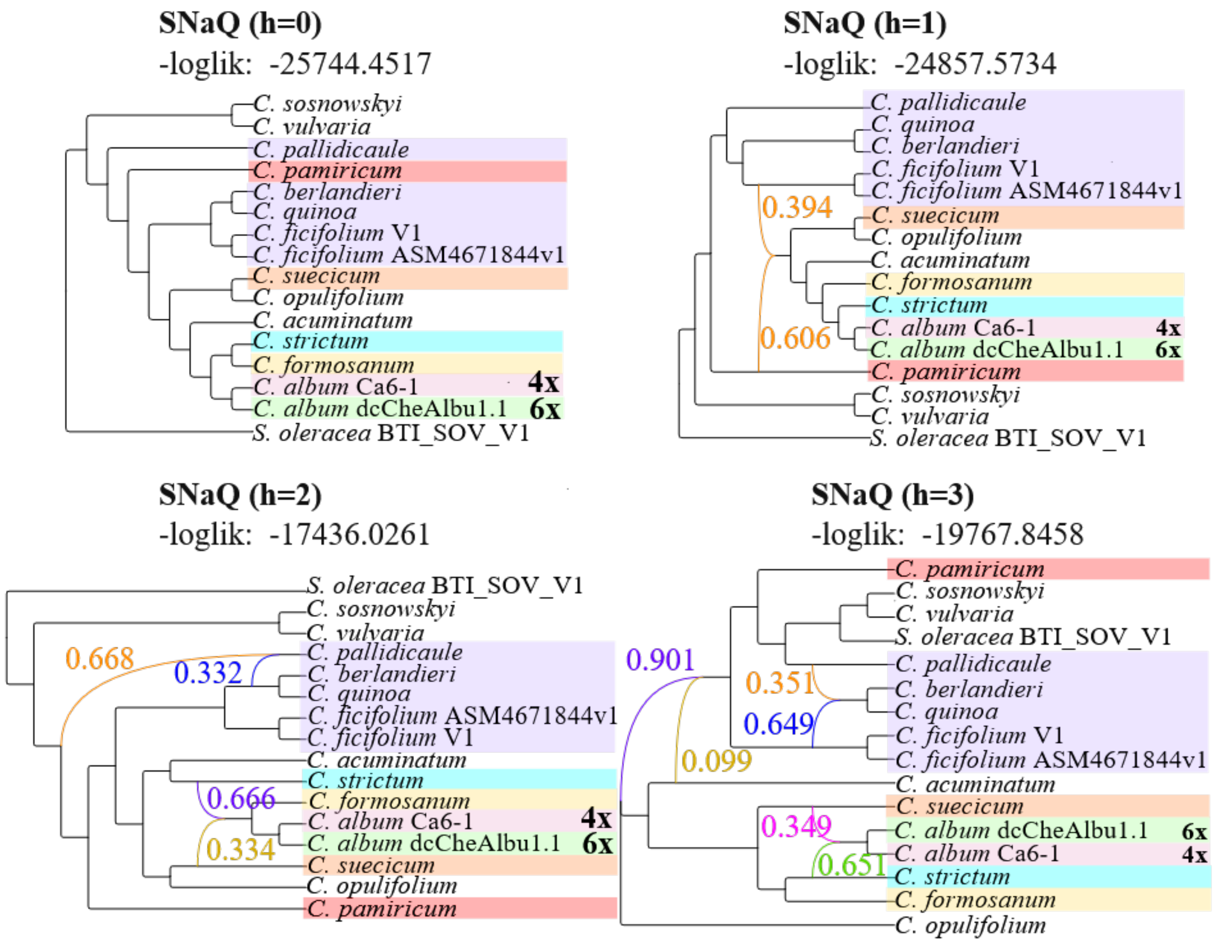


**Supplementary Figure S12. Comparison of phylogenetic network inference using PhyloNet and SNaQ under increasing numbers of reticulations (continued).** (B) SNaQ networks inferred with reticulation numbers ranging from $h = 0$ to $h = 3$. Log-likelihood values are shown for each network. Coloured edges indicate reticulation, where each pair of edges, in complementary colours, sums up to 1. Similar major clades and edge placements were recovered.

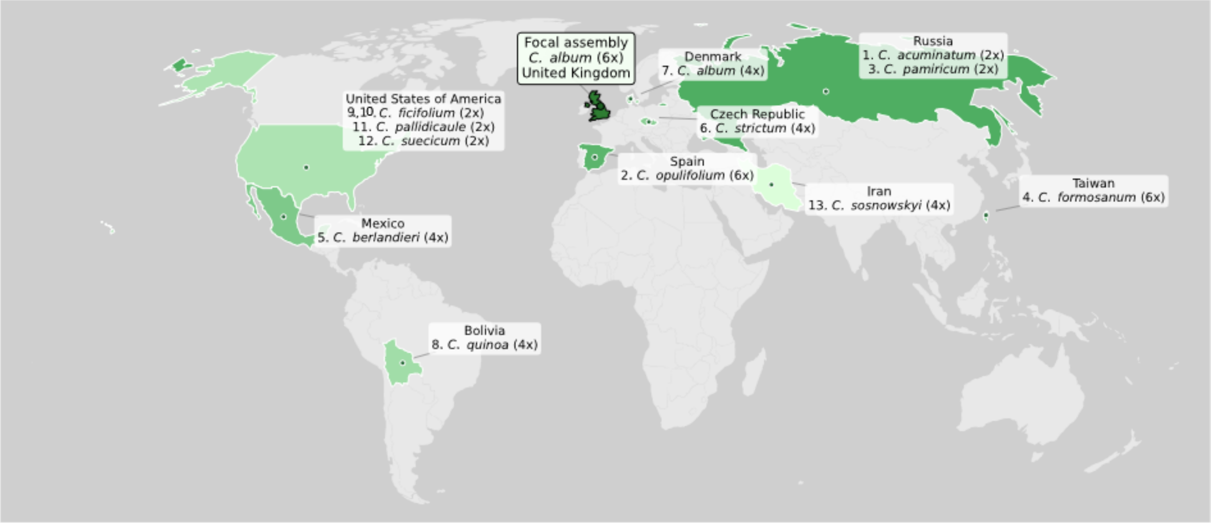


**Supplementary Figure S13. Global distribution of *Chenopodium* genomes contributing to the *C. album* reference assembly dcCheAlbu1.1.** Countries are shaded according to relative genomic contribution, with darker green indicating stronger inferred contributors. Numbers correspond to the ranked contribution of each species to the focal *C. album* genome assembly. The focal assembly (*C. album* dcCheAlbu1.1) originating from the United Kingdom is highlighted. Species names are shown with their ploidy levels in parentheses.

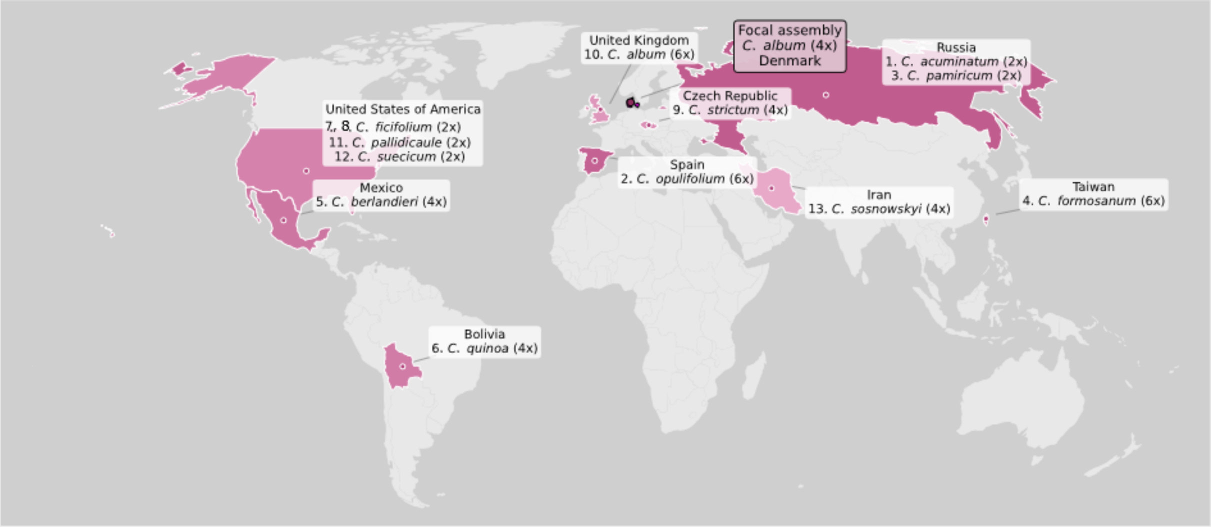


**Supplementary Figure S14. Global distribution of *Chenopodium* genomes contributing to the *C. album* reference assembly Ca6-1.** Countries are shaded according to relative genomic contribution, with darker pink indicating stronger inferred contributors. Numbers correspond to the ranked contribution of each species to the focal *C. album* genome assembly. The focal assembly (*C. album* Ca6-1) originating from the Denmark is highlighted. Species names are shown with their ploidy levels in parentheses.

**Supplementary Table S8. BUSCO loci identified as introgressed in both *C. album* cytotypes.**

| BUSCO ID | Gene | Link |
|---|---|---|
| 100703at71240 | Aspartate/glutamate/uridylate kinase | https://v10-1.orthodb.org/?query=100703at71240 |
| 101246at71240 | Ribosomal RNA small subunit methyltransferase H | https://v10-1.orthodb.org/?query=101246at71240 |
| 10132at71240 | Gamma-tubulin complex component | https://v10-1.orthodb.org/?query=10132at71240 |
| 104541at71240 | AP2/ERF domain | https://v10-1.orthodb.org/?query=104541at71240 |
| 104876at71240 | Hemimethylated DNA-binding domain | https://v10-1.orthodb.org/?query=104876at71240 |
| 105452at71240 | uncharacterized protein LOC107878315 | https://v10-1.orthodb.org/?query=105452at71240 |
| 10648at71240 | Phosphatidylinositol 3-/4-kinase catalytic domain | https://v10-1.orthodb.org/?query=10648at71240 |
| 108717at71240 | Adenosine deaminase/editase | https://v10-1.orthodb.org/?query=108717at71240 |
| 109691at71240 | Translin | https://v10-1.orthodb.org/?query=109691at71240 |
| 110321at71240 | Bromodomain | https://v10-1.orthodb.org/?query=110321at71240 |
| 11057at71240 | Ankyrin repeat-containing domain | https://v10-1.orthodb.org/?query=11057at71240 |

*Continued on next page*

**Supplementary Table S8. BUSCO loci identified as introgressed in both *C. album* cytotypes.**

| BUSCO ID | Gene | Link |
|---|---|---|
| 110600at71240 | Nucleoporin Nup54 alpha-helical domain | https://v10-1.orthodb.org/?query=110600at71240 |
| 112199at71240 | Vps72/YL1 C-terminal | https://v10-1.orthodb.org/?query=112199at71240 |
| 11226at71240 | ATPase AAA-type conserved site | https://v10-1.orthodb.org/?query=11226at71240 |
| 11229at71240 | UvrD-like DNA helicase C-terminal | https://v10-1.orthodb.org/?query=11229at71240 |
| 11295at71240 | Conserved oligomeric Golgi complex subunit 7 | https://v10-1.orthodb.org/?query=11295at71240 |
| 114453at71240 | Spindle and kinetochore-associated protein 1 | https://v10-1.orthodb.org/?query=114453at71240 |
| 115480at71240 | SNARE associated Golgi protein | https://v10-1.orthodb.org/?query=115480at71240 |
| 11773at71240 | Protein TIC110 chloroplastic | https://v10-1.orthodb.org/?query=11773at71240 |
| 119251at71240 | tRNA (guanine-N-7) methyltransferase Trmb type | https://v10-1.orthodb.org/?query=119251at71240 |
| 120257at71240 | CGLD27-like | https://v10-1.orthodb.org/?query=120257at71240 |
| 120543at71240 | Major intrinsic protein | https://v10-1.orthodb.org/?query=120543at71240 |

*Continued on next page*

**Supplementary Table S8. BUSCO loci identified as introgressed in both *C. album* cytotypes.**

| BUSCO ID | Gene | Link |
|---|---|---|
| 121392at71240 | Mucin-like protein | https://v10-1.orthodb.org/?query=121392at71240 |
| 121795at71240 | Phosphoribosylformyl-glycinamidine synthase | https://v10-1.orthodb.org/?query=121795at71240 |
| 12263at71240 | ATPase AAA-type conserved site | https://v10-1.orthodb.org/?query=12263at71240 |
| 12504at71240 | Putative zinc-RING and/or ribbon | https://v10-1.orthodb.org/?query=12504at71240 |
| 126268at71240 | Translation initiation factor 3 | https://v10-1.orthodb.org/?query=126268at71240 |
| 12925at71240 | DNA mismatch repair protein MutS core | https://v10-1.orthodb.org/?query=12925at71240 |
| 129496at71240 | uncharacterized protein LOC101498343 | https://v10-1.orthodb.org/?query=129496at71240 |
| 129792at71240 | RNA-binding S4 domain | https://v10-1.orthodb.org/?query=129792at71240 |
| 131319at71240 | Ribosome-associated YjgA | https://v10-1.orthodb.org/?query=131319at71240 |
| 13273at71240 | Pentatricopeptide repeat | https://v10-1.orthodb.org/?query=13273at71240 |
| 13359at71240 | Armadillo-type fold | https://v10-1.orthodb.org/?query=13359at71240 |

*Continued on next page*

**Supplementary Table S8. BUSCO loci identified as introgressed in both *C. album* cytotypes.**

| BUSCO ID | Gene | Link |
|---|---|---|
| 134282at71240 | PsbP C-terminal | https://v10-1.orthodb.org/?query=134282at71240 |
| 1357at71240 | WD40-repeat-containing domain | https://v10-1.orthodb.org/?query=1357at71240 |
| 137644at71240 | Pinin/SDK/MemA protein | https://v10-1.orthodb.org/?query=137644at71240 |
| 138178at71240 | Nucleoporin NSP1-like C-terminal | https://v10-1.orthodb.org/?query=138178at71240 |
| 138740at71240 | Ribonuclease H superfamily | https://v10-1.orthodb.org/?query=138740at71240 |
| 13917at71240 | BTB/POZ domain | https://v10-1.orthodb.org/?query=13917at71240 |
| 140207at71240 | CTLH C-terminal LisH motif | https://v10-1.orthodb.org/?query=140207at71240 |
| 140209at71240 | Meiotic nuclear division protein 1 homolog | https://v10-1.orthodb.org/?query=140209at71240 |
| 140907at71240 | Pentatricopeptide repeat | https://v10-1.orthodb.org/?query=140907at71240 |
| 141481at71240 | uncharacterized protein LOC107802661 | https://v10-1.orthodb.org/?query=141481at71240 |
| 142714at71240 | Ankyrin repeat-containing domain | https://v10-1.orthodb.org/?query=142714at71240 |

*Continued on next page*

**Supplementary Table S8. BUSCO loci identified as introgressed in both *C. album* cytotypes.**

| BUSCO ID | Gene | Link |
|---|---|---|
| 14405at71240 | DNA2/NAM7 helicase AAA domain | https://v10-1.orthodb.org/?query=14405at71240 |
| 14440at71240 | Tubulin–tyrosine ligase-like protein 12 | https://v10-1.orthodb.org/?query=14440at71240 |
| 144880at71240 | HIT-like domain | https://v10-1.orthodb.org/?query=144880at71240 |
| 145400at71240 | CS domain | https://v10-1.orthodb.org/?query=145400at71240 |
| 146214at71240 | DNA repair protein XRCC4 | https://v10-1.orthodb.org/?query=146214at71240 |
| 147042at71240 | PLAC8 motif-containing protein | https://v10-1.orthodb.org/?query=147042at71240 |
| 147267at71240 | Dimethylallyl adenosine tRNA methylthiotransferase | https://v10-1.orthodb.org/?query=147267at71240 |
| 147278at71240 | DnaJ domain | https://v10-1.orthodb.org/?query=147278at71240 |
| 14824at71240 | DNA mismatch repair protein MutS core | https://v10-1.orthodb.org/?query=14824at71240 |
| 14960at71240 | Pentatricopeptide repeat | https://v10-1.orthodb.org/?query=14960at71240 |
| 14973at71240 | Pentatricopeptide repeat | https://v10-1.orthodb.org/?query=14973at71240 |

*Continued on next page*

**Supplementary Table S8. BUSCO loci identified as introgressed in both *C. album* cytotypes.**

| BUSCO ID | Gene | Link |
|---|---|---|
| 151089at71240 | Peptidyl-prolyl cis-trans isomerase | https://v10-1.orthodb.org/?query=151089at71240 |
| 153075at71240 | Domain of unknown function DUF304 | https://v10-1.orthodb.org/?query=153075at71240 |
| 154419at71240 | Putative pre-16S rRNA nuclease | https://v10-1.orthodb.org/?query=154419at71240 |
| 15628at71240 | HRDC domain | https://v10-1.orthodb.org/?query=15628at71240 |
| 156724at71240 | Rossmann-like alpha/beta/alpha sandwich fold | https://v10-1.orthodb.org/?query=156724at71240 |
| 1578at71240 | SNF2-related N-terminal domain | https://v10-1.orthodb.org/?query=1578at71240 |
| 15904at71240 | Dynamin superfamily | https://v10-1.orthodb.org/?query=15904at71240 |
| 1604at71240 | ATPase AAA-type conserved site | https://v10-1.orthodb.org/?query=1604at71240 |
| 16511at71240 | GPI ethanolamine phosphate transferase 2 isoform X1 | https://v10-1.orthodb.org/?query=16511at71240 |
| 165282at71240 | pistil-specific extensin-like protein | https://v10-1.orthodb.org/?query=165282at71240 |
| 167140at71240 | UDP-N-acetylmuramoyl-L-alanyl-D-glutamate-2 6-diaminopimelate ligase | https://v10-1.orthodb.org/?query=167140at71240 |

*Continued on next page*

**Supplementary Table S8. BUSCO loci identified as introgressed in both *C. album* cytotypes.**

| BUSCO ID | Gene | Link |
|---|---|---|
| 1674at71240 | Nuclear pore complex protein NUP160 | https://v10-1.orthodb.org/?query=1674at71240 |
| 167757at71240 | uncharacterized protein LOC107861211 | https://v10-1.orthodb.org/?query=167757at71240 |
| 17637at71240 | Zinc finger SWIM-type | https://v10-1.orthodb.org/?query=17637at71240 |
| 18261at71240 | Pyridoxal phosphate-dependent transferase | https://v10-1.orthodb.org/?query=18261at71240 |
| 19004at71240 | Protein of unknown function DUF3754 | https://v10-1.orthodb.org/?query=19004at71240 |
| 19271at71240 | Gamma tubulin complex component C-terminal | https://v10-1.orthodb.org/?query=19271at71240 |
| 1950at71240 | Dicer dimerisation domain | https://v10-1.orthodb.org/?query=1950at71240 |
| 19833at71240 | Chaperone J-domain superfamily | https://v10-1.orthodb.org/?query=19833at71240 |
| 19871at71240 | ABC transporter-like | https://v10-1.orthodb.org/?query=19871at71240 |
| 2018at71240 | WD40 repeat | https://v10-1.orthodb.org/?query=2018at71240 |
| 204at71240 | Translational activator Gcn1 | https://v10-1.orthodb.org/?query=204at71240 |
| 20830at71240 | RNA-binding CRM domain | https://v10-1.orthodb.org/?query=20830at71240 |

*Continued on next page*

**Supplementary Table S8. BUSCO loci identified as introgressed in both *C. album* cytotypes.**

| BUSCO ID | Gene | Link |
|---|---|---|
| 20867at71240 | Ku C-terminal | https://v10-1.orthodb.org/?query=20867at71240 |
| 21873at71240 | Anaphase-promoting complex subunit 4 long domain | https://v10-1.orthodb.org/?query=21873at71240 |
| 21992at71240 | FG-GAP repeat-containing protein | https://v10-1.orthodb.org/?query=21992at71240 |
| 22403at71240 | Tetratricopeptide repeat | https://v10-1.orthodb.org/?query=22403at71240 |
| 224at71240 | WD40 repeat | https://v10-1.orthodb.org/?query=224at71240 |
| 22952at71240 | Beta-galactosidase | https://v10-1.orthodb.org/?query=22952at71240 |
| 22981at71240 | Timeless protein | https://v10-1.orthodb.org/?query=22981at71240 |
| 23003at71240 | GPI transamidase component Gaa1 | https://v10-1.orthodb.org/?query=23003at71240 |
| 23032at71240 | DNA mismatch repair protein Mlh1 C-terminal | https://v10-1.orthodb.org/?query=23032at71240 |
| 23089at71240 | ABC transporter type 1 transmembrane domain | https://v10-1.orthodb.org/?query=23089at71240 |
| 2327at71240 | Polymerase nucleotidyl transferase domain | https://v10-1.orthodb.org/?query=2327at71240 |
| 23463at71240 | SANT/Myb domain | https://v10-1.orthodb.org/?query=23463at71240 |

*Continued on next page*

**Supplementary Table S8. BUSCO loci identified as introgressed in both *C. album* cytotypes.**

| BUSCO ID | Gene | Link |
|---|---|---|
| 23535at71240 | DNA polymerase Y-family little finger domain | https://v10-1.orthodb.org/?query=23535at71240 |
| 23684at71240 | Nucleic acid-binding OB-fold | https://v10-1.orthodb.org/?query=23684at71240 |
| 2395at71240 | WD40 repeat | https://v10-1.orthodb.org/?query=2395at71240 |
| 24945at71240 | Pentatricopeptide repeat | https://v10-1.orthodb.org/?query=24945at71240 |
| 25047at71240 | Protein kinase domain | https://v10-1.orthodb.org/?query=25047at71240 |
| 25476at71240 | MoeA C-terminal domain IV | https://v10-1.orthodb.org/?query=25476at71240 |
| 26288at71240 | Endonuclease/exonuclease/ phosphatase | https://v10-1.orthodb.org/?query=26288at71240 |
| 27496at71240 | Leucine-rich repeat | https://v10-1.orthodb.org/?query=27496at71240 |
| 28331at71240 | Pentatricopeptide repeat | https://v10-1.orthodb.org/?query=28331at71240 |
| 2884at71240 | Tetratricopeptide repeat | https://v10-1.orthodb.org/?query=2884at71240 |
| 28907at71240 | Calcium-dependent channel 7TM region putative phosphate | https://v10-1.orthodb.org/?query=28907at71240 |

*Continued on next page*

**Supplementary Table S8. BUSCO loci identified as introgressed in both *C. album* cytotypes.**

| BUSCO ID | Gene | Link |
|---|---|---|
| 29664at71240 | Alpha/Beta hydrolase fold | https://v10-1.orthodb.org/?query=29664at71240 |
| 29755at71240 | Anoctamin | https://v10-1.orthodb.org/?query=29755at71240 |
| 29842at71240 | NAD(P)-binding domain | https://v10-1.orthodb.org/?query=29842at71240 |
| 30785at71240 | Nucleolar complex protein 2 | https://v10-1.orthodb.org/?query=30785at71240 |
| 30832at71240 | WD40-repeat-containing domain | https://v10-1.orthodb.org/?query=30832at71240 |
| 31816at71240 | Helicase C-terminal | https://v10-1.orthodb.org/?query=31816at71240 |
| 32201at71240 | Helicase C-terminal | https://v10-1.orthodb.org/?query=32201at71240 |
| 32378at71240 | alpha-12-Mannosidase | https://v10-1.orthodb.org/?query=32378at71240 |
| 32626at71240 | GDP-fucose protein O-fucosyltransferase | https://v10-1.orthodb.org/?query=32626at71240 |
| 32788at71240 | Starch synthase catalytic domain | https://v10-1.orthodb.org/?query=32788at71240 |
| 33442at71240 | Subtilisin-like protease fibronectin type-III domain | https://v10-1.orthodb.org/?query=33442at71240 |

*Continued on next page*

**Supplementary Table S8. BUSCO loci identified as introgressed in both *C. album* cytotypes.**

| BUSCO ID | Gene | Link |
|---|---|---|
| 33746at71240 | GDP-fucose protein O-fucosyltransferase | https://v10-1.orthodb.org/?query=33746at71240 |
| 3391at71240 | TATA-binding protein interacting (TIP20) | https://v10-1.orthodb.org/?query=3391at71240 |
| 34656at71240 | Nuclear fragile X mental retardation-interacting protein 1 conserved domain | https://v10-1.orthodb.org/?query=34656at71240 |
| 34798at71240 | Transglutaminase-like | https://v10-1.orthodb.org/?query=34798at71240 |
| 35316at71240 | tRNA (guanine(37)-N1)-methyltransferase | https://v10-1.orthodb.org/?query=35316at71240 |
| 36003at71240 | Ankyrin repeat-containing domain | https://v10-1.orthodb.org/?query=36003at71240 |
| 36040at71240 | Nitroreductase | https://v10-1.orthodb.org/?query=36040at71240 |
| 36224at71240 | Cwf19-like C-terminal domain-1 | https://v10-1.orthodb.org/?query=36224at71240 |
| 36501at71240 | UV-stimulated scaffold protein A | https://v10-1.orthodb.org/?query=36501at71240 |
| 368at71240 | Armadillo-type fold | https://v10-1.orthodb.org/?query=368at71240 |
| 39529at71240 | Thiamine pyrophosphate enzyme C-terminal TPP-binding | https://v10-1.orthodb.org/?query=39529at71240 |
| 39696at71240 | Forkhead-associated (FHA) domain | https://v10-1.orthodb.org/?query=39696at71240 |

*Continued on next page*

**Supplementary Table S8. BUSCO loci identified as introgressed in both *C. album* cytotypes.**

| BUSCO ID | Gene | Link |
|---|---|---|
| 40417at71240 | MORN motif | https://v10-1.orthodb.org/?query=40417at71240 |
| 4043at71240 | SH3 domain | https://v10-1.orthodb.org/?query=4043at71240 |
| 40667at71240 | CASP C-terminal | https://v10-1.orthodb.org/?query=40667at71240 |
| 41496at71240 | Sec1-like protein | https://v10-1.orthodb.org/?query=41496at71240 |
| 41635at71240 | K Homology domain type 1 | https://v10-1.orthodb.org/?query=41635at71240 |
| 41819at71240 | Light-mediated development protein DET1 | https://v10-1.orthodb.org/?query=41819at71240 |
| 42397at71240 | Carbohydrate kinase FGGY C-terminal | https://v10-1.orthodb.org/?query=42397at71240 |
| 42494at71240 | Pectinesterase | https://v10-1.orthodb.org/?query=42494at71240 |
| 42509at71240 | Mu homology domain | https://v10-1.orthodb.org/?query=42509at71240 |
| 43003at71240 | Peroxisomal/glyoxysomal leader peptide-processing protease | https://v10-1.orthodb.org/?query=43003at71240 |
| 4325at71240 | Helicase C-terminal | https://v10-1.orthodb.org/?query=4325at71240 |

*Continued on next page*

**Supplementary Table S8. BUSCO loci identified as introgressed in both *C. album* cytotypes.**

| BUSCO ID | Gene | Link |
|---|---|---|
| 43419at71240 | paramyosin | https://v10-1.orthodb.org/?query=43419at71240 |
| 43815at71240 | GTP binding domain | https://v10-1.orthodb.org/?query=43815at71240 |
| 4403at71240 | tRNA wybutosine-synthesizing protein | https://v10-1.orthodb.org/?query=4403at71240 |
| 45115at71240 | Enolase | https://v10-1.orthodb.org/?query=45115at71240 |
| 45155at71240 | Glycosyltransferase 61 | https://v10-1.orthodb.org/?query=45155at71240 |
| 45216at71240 | uncharacterized protein LOC101502623 | https://v10-1.orthodb.org/?query=45216at71240 |
| 45959at71240 | Isopenicillin N synthase-like | https://v10-1.orthodb.org/?query=45959at71240 |
| 46667at71240 | Mini-chromosome maintenance complex-binding protein | https://v10-1.orthodb.org/?query=46667at71240 |
| 46995at71240 | Diacylglycerol kinase catalytic domain | https://v10-1.orthodb.org/?query=46995at71240 |
| 47260at71240 | Histidine phosphatase superfamily | https://v10-1.orthodb.org/?query=47260at71240 |
| 4807at71240 | Nrap protein domain 4 | https://v10-1.orthodb.org/?query=4807at71240 |

*Continued on next page*

**Supplementary Table S8. BUSCO loci identified as introgressed in both *C. album* cytotypes.**

| BUSCO ID | Gene | Link |
|---|---|---|
| 48163at71240 | Leucine-rich repeat | https://v10-1.orthodb.org/?query=48163at71240 |
| 48330at71240 | Kelch repeat type 1 | https://v10-1.orthodb.org/?query=48330at71240 |
| 48817at71240 | MIP18 family-like | https://v10-1.orthodb.org/?query=48817at71240 |
| 49259at71240 | Peptidase M24 | https://v10-1.orthodb.org/?query=49259at71240 |
| 50768at71240 | GPI mannosyltransferase 2 | https://v10-1.orthodb.org/?query=50768at71240 |
| 50811at71240 | NAD(P)-binding domain | https://v10-1.orthodb.org/?query=50811at71240 |
| 51218at71240 | golgin candidate 2 | https://v10-1.orthodb.org/?query=51218at71240 |
| 51378at71240 | Pseudouridine synthase catalytic domain superfamily | https://v10-1.orthodb.org/?query=51378at71240 |
| 526at71240 | ABC transporter A | https://v10-1.orthodb.org/?query=526at71240 |
| 52731at71240 | Ankyrin repeat-containing domain | https://v10-1.orthodb.org/?query=52731at71240 |
| 53606at71240 | Major facilitator superfamily | https://v10-1.orthodb.org/?query=53606at71240 |
| 5406at71240 | SecA preprotein cross-linking domain | https://v10-1.orthodb.org/?query=5406at71240 |

*Continued on next page*

**Supplementary Table S8. BUSCO loci identified as introgressed in both *C. album* cytotypes.**

| BUSCO ID | Gene | Link |
|---|---|---|
| 55525at71240 | Ribosomal protein Rsm22-like | https://v10-1.orthodb.org/?query=55525at71240 |
| 55986at71240 | Zinc finger PHD-type | https://v10-1.orthodb.org/?query=55986at71240 |
| 56709at71240 | uncharacterized protein LOC107816884 | https://v10-1.orthodb.org/?query=56709at71240 |
| 56817at71240 | Mannosyltransferase | https://v10-1.orthodb.org/?query=56817at71240 |
| 58086at71240 | Metallo-beta-lactamase | https://v10-1.orthodb.org/?query=58086at71240 |
| 59794at71240 | VASt domain | https://v10-1.orthodb.org/?query=59794at71240 |
| 597at71240 | Dicer dimerisation domain | https://v10-1.orthodb.org/?query=597at71240 |
| 60063at71240 | RED-like N-terminal | https://v10-1.orthodb.org/?query=60063at71240 |
| 61505at71240 | Histone acetyltransferase type B catalytic subunit | https://v10-1.orthodb.org/?query=61505at71240 |
| 62739at71240 | Dolichol kinase | https://v10-1.orthodb.org/?query=62739at71240 |
| 62741at71240 | Reticulon | https://v10-1.orthodb.org/?query=62741at71240 |
| 62825at71240 | Peptidase M24 | https://v10-1.orthodb.org/?query=62825at71240 |

*Continued on next page*

**Supplementary Table S8. BUSCO loci identified as introgressed in both *C. album* cytotypes.**

| BUSCO ID | Gene | Link |
|---|---|---|
| 62949at71240 | 3-deoxy-D-manno-octulosonic-acid transferase N-terminal | https://v10-1.orthodb.org/?query=62949at71240 |
| 63750at71240 | Malonyl-CoA decarboxylase | https://v10-1.orthodb.org/?query=63750at71240 |
| 64632at71240 | Actin family | https://v10-1.orthodb.org/?query=64632at71240 |
| 64680at71240 | WD40 repeat | https://v10-1.orthodb.org/?query=64680at71240 |
| 66230at71240 | DNA primase large subunit | https://v10-1.orthodb.org/?query=66230at71240 |
| 66590at71240 | GTP binding domain | https://v10-1.orthodb.org/?query=66590at71240 |
| 6689at71240 | Alpha-16-glucosidases pullulanase-type | https://v10-1.orthodb.org/?query=6689at71240 |
| 68088at71240 | Rad9/Ddc1 | https://v10-1.orthodb.org/?query=68088at71240 |
| 68100at71240 | Clathrin adaptor mu subunit | https://v10-1.orthodb.org/?query=68100at71240 |
| 68407at71240 | WW domain | https://v10-1.orthodb.org/?query=68407at71240 |
| 68776at71240 | Beta-ketoacyl synthase C-terminal | https://v10-1.orthodb.org/?query=68776at71240 |

*Continued on next page*

**Supplementary Table S8. BUSCO loci identified as introgressed in both *C. album* cytotypes.**

| BUSCO ID | Gene | Link |
|---|---|---|
| 6890at71240 | Pentatricopeptide repeat | https://v10-1.orthodb.org/?query=6890at71240 |
| 69753at71240 | Methyltransferase domain | https://v10-1.orthodb.org/?query=69753at71240 |
| 70311at71240 | Cyclophilin-type peptidyl-prolyl cis-trans isomerase domain | https://v10-1.orthodb.org/?query=70311at71240 |
| 70527at71240 | Glycosyl transferase family 19 | https://v10-1.orthodb.org/?query=70527at71240 |
| 71100at71240 | Embryo defective 2737 | https://v10-1.orthodb.org/?query=71100at71240 |
| 71626at71240 | Tsl-kinase interacting protein 1 | https://v10-1.orthodb.org/?query=71626at71240 |
| 72494at71240 | Pentatricopeptide repeat | https://v10-1.orthodb.org/?query=72494at71240 |
| 74810at71240 | TraB family | https://v10-1.orthodb.org/?query=74810at71240 |
| 75421at71240 | Amino acid/polyamine transporter I | https://v10-1.orthodb.org/?query=75421at71240 |
| 75656at71240 | Negative regulator of systemic acquired resistance SNI1 | https://v10-1.orthodb.org/?query=75656at71240 |
| 75917at71240 | Las1 | https://v10-1.orthodb.org/?query=75917at71240 |

*Continued on next page*

**Supplementary Table S8. BUSCO loci identified as introgressed in both *C. album* cytotypes.**

| BUSCO ID | Gene | Link |
|---|---|---|
| 75984at71240 | Phosphatidylserine decarboxylase proenzyme 1 mitochondrial | https://v10-1.orthodb.org/?query=75984at71240 |
| 76918at71240 | Probable tRNA N6-adenosine threonylcarbamoyltransferase mitochondrial | https://v10-1.orthodb.org/?query=76918at71240 |
| 77821at71240 | WD40-repeat-containing domain | https://v10-1.orthodb.org/?query=77821at71240 |
| 78140at71240 | Cysteine peptidase cysteine active site | https://v10-1.orthodb.org/?query=78140at71240 |
| 78143at71240 | Regulator of chromosome condensation 1/beta-lactamase-inhibitor protein II | https://v10-1.orthodb.org/?query=78143at71240 |
| 7892at71240 | Importin-beta N-terminal domain | https://v10-1.orthodb.org/?query=7892at71240 |
| 79006at71240 | Leucine-rich repeat | https://v10-1.orthodb.org/?query=79006at71240 |
| 8000at71240 | Armadillo-type fold | https://v10-1.orthodb.org/?query=8000at71240 |
| 80484at71240 | Signal recognition particle SRP54 helical bundle | https://v10-1.orthodb.org/?query=80484at71240 |
| 80865at71240 | Tetratricopeptide-like helical domain superfamily | https://v10-1.orthodb.org/?query=80865at71240 |
| 8091at71240 | AP-5 complex subunit beta-1 | https://v10-1.orthodb.org/?query=8091at71240 |

*Continued on next page*

**Supplementary Table S8. BUSCO loci identified as introgressed in both *C. album* cytotypes.**

| BUSCO ID | Gene | Link |
|---|---|---|
| 81297at71240 | Zinc finger RING-type | https://v10-1.orthodb.org/?query=81297at71240 |
| 82332at71240 | Carbohydrate kinase PfkB | https://v10-1.orthodb.org/?query=82332at71240 |
| 82479at71240 | Domain of unknown function DUF1995 | https://v10-1.orthodb.org/?query=82479at71240 |
| 82679at71240 | F-box domain | https://v10-1.orthodb.org/?query=82679at71240 |
| 82761at71240 | Dual-specificity RNA methyl-transferase RlmN | https://v10-1.orthodb.org/?query=82761at71240 |
| 83256at71240 | NAD-dependent epimerase/dehydratase | https://v10-1.orthodb.org/?query=83256at71240 |
| 84628at71240 | NADP-dependent oxidoreductase domain | https://v10-1.orthodb.org/?query=84628at71240 |
| 84814at71240 | WD40 repeat | https://v10-1.orthodb.org/?query=84814at71240 |
| 86440at71240 | Pseudouridine synthase RsuA/RluA | https://v10-1.orthodb.org/?query=86440at71240 |
| 8739at71240 | DNA mismatch repair protein MutS core | https://v10-1.orthodb.org/?query=8739at71240 |
| 87734at71240 | Root UVB sensitive family | https://v10-1.orthodb.org/?query=87734at71240 |

*Continued on next page*

**Supplementary Table S8. BUSCO loci identified as introgressed in both *C. album* cytotypes.**

| BUSCO ID | Gene | Link |
|---|---|---|
| 87765at71240 | Vacuolar protein sorting-associated protein Ist1 | https://v10-1.orthodb.org/?query=87765at71240 |
| 88382at71240 | BRO1 domain | https://v10-1.orthodb.org/?query=88382at71240 |
| 90287at71240 | Elongator complex protein 5 | https://v10-1.orthodb.org/?query=90287at71240 |
| 90893at71240 | Tryptophan synthase beta subunit-like PLP-dependent enzyme | https://v10-1.orthodb.org/?query=90893at71240 |
| 91349at71240 | Alpha/Beta hydrolase fold | https://v10-1.orthodb.org/?query=91349at71240 |
| 91582at71240 | Chalcone isomerase | https://v10-1.orthodb.org/?query=91582at71240 |
| 91944at71240 | CobW/HypB/UreG nucleotide-binding domain | https://v10-1.orthodb.org/?query=91944at71240 |
| 92777at71240 | Alpha/Beta hydrolase fold | https://v10-1.orthodb.org/?query=92777at71240 |
| 933at71240 | UDP-glucose:Glycoprotein Glucosyltransferase | https://v10-1.orthodb.org/?query=933at71240 |
| 94348at71240 | Leucine carboxyl methyltransferase 1 | https://v10-1.orthodb.org/?query=94348at71240 |
| 94553at71240 | Ubiquitin domain | https://v10-1.orthodb.org/?query=94553at71240 |
| 95643at71240 | PPM-type phosphatase domain | https://v10-1.orthodb.org/?query=95643at71240 |

*Continued on next page*

**Supplementary Table S8. BUSCO loci identified as introgressed in both *C. album* cytotypes.**

| BUSCO ID | Gene | Link |
|---|---|---|
| 96055at71240 | DNA endonuclease Ctp1 C-terminal | https://v10-1.orthodb.org/?query=96055at71240 |
| 9729at71240 | WRC domain | https://v10-1.orthodb.org/?query=9729at71240 |
| 97786at71240 | Plant organelle RNA recognition domain | https://v10-1.orthodb.org/?query=97786at71240 |
| 98387at71240 | Peptide chain release factor | https://v10-1.orthodb.org/?query=98387at71240 |
| 99040at71240 | Leucine-rich repeat cysteine-containing subtype | https://v10-1.orthodb.org/?query=99040at71240 |
| 99284at71240 | Tetratricopeptide-like helical domain superfamily | https://v10-1.orthodb.org/?query=99284at71240 |

**Supplementary Table S9. BUSCO loci identified as introgressed only in *C. album* Ca6-1 4x.**

| BUSCO ID | Gene | Link |
|---|---|---|
| 100163at71240 | S1/P1 nuclease | https://v10-1.orthodb.org/?query=100163at71240 |
| 103322at71240 | Zinc finger CCCH-type | https://v10-1.orthodb.org/?query=103322at71240 |
| 103378at71240 | MEMO1 family | https://v10-1.orthodb.org/?query=103378at71240 |
| 10618at71240 | Gamma-tubulin complex component | https://v10-1.orthodb.org/?query=10618at71240 |

*Continued on next page*

**Supplementary Table S9. BUSCO loci identified as introgressed only in *C. album* Ca6-1 4x.**

| BUSCO ID | Gene | Link |
| --- | --- | --- |
| 106602at71240 | Phosphoribulokinase/uridine kinase | https://v10-1.orthodb.org/?query=106602at71240 |
| 106984at71240 | Tetraketide alpha-pyrone reductase 1 | https://v10-1.orthodb.org/?query=106984at71240 |
| 10881at71240 | Oberon coiled-coil region | https://v10-1.orthodb.org/?query=10881at71240 |
| 110127at71240 | Leucine-rich repeat | https://v10-1.orthodb.org/?query=110127at71240 |
| 110565at71240 | Uncharacterized protein | https://v10-1.orthodb.org/?query=110565at71240 |
| 111827at71240 | Uncharacterized protein | https://v10-1.orthodb.org/?query=111827at71240 |
| 118301at71240 | Magnesium-protoporphyrin IX methyltransferase | https://v10-1.orthodb.org/?query=118301at71240 |
| 121792at71240 | Lysine methyltransferase | https://v10-1.orthodb.org/?query=121792at71240 |
| 1308at71240 | DNA2/NAM7 helicase AAA domain | https://v10-1.orthodb.org/?query=1308at71240 |
| 13125at71240 | Pentatricopeptide repeat (PPR) | https://v10-1.orthodb.org/?query=13125at71240 |
| 139689at71240 | Methyltransferase domain | https://v10-1.orthodb.org/?query=139689at71240 |

*Continued on next page*

**Supplementary Table S9. BUSCO loci identified as introgressed only in *C. album* Ca6-1 4x.**

| BUSCO ID | Gene | Link |
|---|---|---|
| 140463at71240 | GNAT domain | https://v10-1.orthodb.org/?query=140463at71240 |
| 1589at71240 | Magnesium-chelatase subunit ChlH | https://v10-1.orthodb.org/?query=1589at71240 |
| 18602at71240 | BAG domain | https://v10-1.orthodb.org/?query=18602at71240 |
| 22901at71240 | Helicase C-terminal | https://v10-1.orthodb.org/?query=22901at71240 |
| 2956at71240 | Leucine-rich repeat | https://v10-1.orthodb.org/?query=2956at71240 |
| 35653at71240 | WD40 repeat | https://v10-1.orthodb.org/?query=35653at71240 |
| 37422at71240 | RNA cap guanine-N2 methyltransferase | https://v10-1.orthodb.org/?query=37422at71240 |
| 40388at71240 | Peptidase C78 (Ulp/SUMO protease) | https://v10-1.orthodb.org/?query=40388at71240 |
| 41252at71240 | Nucleolar protein 14 | https://v10-1.orthodb.org/?query=41252at71240 |
| 41443at71240 | Protein Lines N-terminal | https://v10-1.orthodb.org/?query=41443at71240 |
| 52900at71240 | Rubisco LSMT substrate-binding domain | https://v10-1.orthodb.org/?query=52900at71240 |

*Continued on next page*

**Supplementary Table S9. BUSCO loci identified as introgressed only in *C. album* Ca6-1 4x.**

| BUSCO ID | Gene | Link |
|---|---|---|
| 5623at71240 | NHL repeat | https://v10-1.orthodb.org/?query=5623at71240 |
| 56887at71240 | SAM-dependent methyltransferase | https://v10-1.orthodb.org/?query=56887at71240 |
| 6058at71240 | ATPase AAA-type | https://v10-1.orthodb.org/?query=6058at71240 |
| 65502at71240 | Pentatricopeptide repeat | https://v10-1.orthodb.org/?query=65502at71240 |
| 65978at71240 | Pentatricopeptide repeat | https://v10-1.orthodb.org/?query=65978at71240 |
| 6688at71240 | Glycoside hydrolase family 77 | https://v10-1.orthodb.org/?query=6688at71240 |
| 86274at71240 | Chloroplast envelope membrane protein CemA | https://v10-1.orthodb.org/?query=86274at71240 |
| 86at71240 | Phosphatidylinositol 3-/4-kinase | https://v10-1.orthodb.org/?query=86at71240 |
| 88282at71240 | WD40 repeat | https://v10-1.orthodb.org/?query=88282at71240 |
| 93922at71240 | AAR2 N-terminal | https://v10-1.orthodb.org/?query=93922at71240 |
| 94424at71240 | Uncharacterized protein AT4g12540 | https://v10-1.orthodb.org/?query=94424at71240 |
| 9527at71240 | N-acetyltransferase B complex subunit | https://v10-1.orthodb.org/?query=9527at71240 |

*Continued on next page*

**Supplementary Table S9. BUSCO loci identified as introgressed only in *C. album* Ca6-1 4x.**

| BUSCO ID | Gene | Link |
|---|---|---|
| 97605at71240 | Carbohydrate kinase PfkB | https://v10-1.orthodb.org/?query=97605at71240 |

**Supplementary Table S10. BUSCO loci identified as introgressed only in *C. album* dcCheAlbu1.1 6x**

| BUSCO ID | Gene | Link |
|---|---|---|
| 104482at71240 | FAD dependent oxidoreductase | https://v10-1.orthodb.org/?query=104482at71240 |
| 109983at71240 | NmrA-like domain | https://v10-1.orthodb.org/?query=109983at71240 |
| 10at71240 | Zinc finger ZZ-type | https://v10-1.orthodb.org/?query=10at71240 |
| 112753at71240 | DDT domain | https://v10-1.orthodb.org/?query=112753at71240 |
| 1159at71240 | Nucleolar 27S pre-rRNA processing | https://v10-1.orthodb.org/?query=1159at71240 |
| 12736at71240 | Urease | https://v10-1.orthodb.org/?query=12736at71240 |
| 134575at71240 | Mog1/PsbP alpha/beta/alpha sandwich | https://v10-1.orthodb.org/?query=134575at71240 |
| 135744at71240 | Zinc finger PHD-type | https://v10-1.orthodb.org/?query=135744at71240 |
| 139365at71240 | Oxygen-evolving enhancer protein 3 | https://v10-1.orthodb.org/?query=139365at71240 |

*Continued on next page*

**Supplementary Table S10. BUSCO loci identified as introgressed only in *C. album* dcCheAlbu1.1 6x**

| BUSCO ID | Gene | Link |
|---|---|---|
| 139835at71240 | DUF4079 protein | https://v10-1.orthodb.org/?query=139835at71240 |
| 15341at71240 | MCM domain | https://v10-1.orthodb.org/?query=15341at71240 |
| 156871at71240 | Ankyrin repeat-containing domain | https://v10-1.orthodb.org/?query=156871at71240 |
| 164300at71240 | Uncharacterized protein | https://v10-1.orthodb.org/?query=164300at71240 |
| 22497at71240 | G patch domain protein | https://v10-1.orthodb.org/?query=22497at71240 |
| 2421at71240 | TRAPP II complex Trs120 | https://v10-1.orthodb.org/?query=2421at71240 |
| 2at71240 | Midasin AAA lid domain | https://v10-1.orthodb.org/?query=2at71240 |
| 312at71240 | Zinc finger FYVE domain | https://v10-1.orthodb.org/?query=312at71240 |
| 35634at71240 | tRNA-dihydrouridine synthase | https://v10-1.orthodb.org/?query=35634at71240 |
| 38328at71240 | DNA-directed primase/polymerase | https://v10-1.orthodb.org/?query=38328at71240 |
| 38450at71240 | Pentatricopeptide repeat | https://v10-1.orthodb.org/?query=38450at71240 |
| 40627at71240 | Aldehyde dehydrogenase | https://v10-1.orthodb.org/?query=40627at71240 |

*Continued on next page*

**Supplementary Table S10. BUSCO loci identified as introgressed only in *C. album* dcCheAlbu1.1 6x**

| BUSCO ID | Gene | Link |
|---|---|---|
| 41693at71240 | AMP-binding conserved site | https://v10-1.orthodb.org/?query=41693at71240 |
| 43005at71240 | Kelch repeat | https://v10-1.orthodb.org/?query=43005at71240 |
| 4302at71240 | AP-3 complex subunit beta | https://v10-1.orthodb.org/?query=4302at71240 |
| 44670at71240 | NAD(P)-binding domain | https://v10-1.orthodb.org/?query=44670at71240 |
| 4838at71240 | MIF4G-like domain | https://v10-1.orthodb.org/?query=4838at71240 |
| 50477at71240 | RNA-binding CRM domain | https://v10-1.orthodb.org/?query=50477at71240 |
| 52721at71240 | YAP-binding/ALF4/Glomulin | https://v10-1.orthodb.org/?query=52721at71240 |
| 54140at71240 | NADPH | https://v10-1.orthodb.org/?query=54140at71240 |
| 55641at71240 | Mur ligase central | https://v10-1.orthodb.org/?query=55641at71240 |
| 57561at71240 | Uncharacterized protein At3g14900 | https://v10-1.orthodb.org/?query=57561at71240 |
| 64803at71240 | Rubisco LSMT domain | https://v10-1.orthodb.org/?query=64803at71240 |

*Continued on next page*

**Supplementary Table S10. BUSCO loci identified as introgressed only in *C. album* dcCheAlbu1.1 6x**

| BUSCO ID | Gene | Link |
|---|---|---|
| 6786at71240 | C2 domain | https://v10-1.orthodb.org/?query=6786at71240 |
| 68711at71240 | Myosin heavy chain-related | https://v10-1.orthodb.org/?query=68711at71240 |
| 7309at71240 | SMC5 | https://v10-1.orthodb.org/?query=7309at71240 |
| 793at71240 | Protein virilizer | https://v10-1.orthodb.org/?query=793at71240 |
| 79897at71240 | Rieske protein | https://v10-1.orthodb.org/?query=79897at71240 |
| 86443at71240 | Protein prenyltransferase alpha | https://v10-1.orthodb.org/?query=86443at71240 |
